# Nitrogen availability drives gene length of dominant prokaryotes and diversity of genes acquiring Nitrogen-species in oceanic systems

**DOI:** 10.1101/2021.01.10.426031

**Authors:** Leon Dlugosch, Anja Poehlein, Bernd Wemheuer, Birgit Pfeiffer, Helge-A. Giebel, Rolf Daniel, Meinhard Simon

**Author notes:** Corresponding author: Meinhard Simon.

## Abstract

Nitrogen (N) is a key element for prokaryotes in the oceans and often limits phytoplankton primary production. An untested option to reduce prokaryotic N-demand under N-limitation is to reduce gene length. Here we show that in the sunlit Atlantic Ocean genes of the prokaryotic microbial communities in the permanently stratified N-limited (sub)tropics are up to 20% shorter than in N-replete regions further south and north. Average gene length (AGL) of major pelagic prokaryotic genera and two virus families correlated positively with nitrate concentrations. Further, the genomic G+C content of 60% of the taxa was lower and the gene repertoire to acquire inorganic and organic N-species higher in N-limited than in N-replete regions. A comparison of the N-demand by reducing gene length or G+C content showed that the former is much more efficient to save N. Our findings introduce a novel and most effective mode of evolutionary adaptation of prokaryotes to save resources including N and energy. They further show an enhanced diversification of genes acquiring N-species and -compounds in N-deplete relative to N-replete regions and thus add important information for a better understanding of the evolutionary adaptation of prokaryotes to N-availability in oceanic systems.

## Main

Genome evolution in prokaryotes is largely driven by mutation and horizontal gene transfer (HGT) resulting in acquisition and deletion of genes^1–3^. Whereas HGT leads to gain or loss of entire genes or gene clusters, mutation initially leads to gene modification, either non-synonymous or synonymous, and possibly to deletion of codons or pseudogenes and eventually of genes^1,4,5^. Gene length is thus affected by mutational changes and reflected in the variation of AGL in different prokaryotes^1^. However, it is unknown whether gene length is affected by evolutionary constraints such as growth limitation by nutrients or elements such as N. Under relaxing growth conditions, evolving prokaryotes increase their genomic G+C content to improve their fitness, driven by mutation bias and other, not fully understood selective forces^3,6^. The genomic G+C content of the majority of auto- and heterotrophic bacterial classes and families is positively correlated with their genome size^7^, implying that genome expansion by acquisition of beneficial traits via HGT increases the G+C content. Despite this general trend, under strong environmental constraints genome size, the genomic G+C content, and purifying selection towards a reduced genomic N-content of prokaryotes underlies selective forces leading to a reduced G+C content and genome size to utilize limiting resources more efficiently. A consistently low ratio of nonsynonymous polymorphisms to synonymous polymorphisms in the Tara Ocean prokaryotic gene set and the identification of nitrate as the strongest environmental variable correlating with this ratio indicates that purifying selection drives the reduced genomic N-content of oceanic prokaryotes^8^. The relatively low G+C content and small genomes of bacterial lineages in stratified oceans, in particular the abundant cyanobacterium *Prochlorococcus*, the alphaproteobacterial *Pelagibacteraceae*/SAR11 and the gammaproteobacterial SAR86 clades, are the result of genome streamlining^2^ and strong N-limitation^4,9^ because the nucleobases G+C require one atom more N than A+T. This strong forcing by N-availability and minimizing N-cost affects also the proteome of these pelagic bacterial lineages^9,10^. Genes with a lower G+C content encode amino acids with reduced N per amino acid residue side chain (N-ARSC) even though the mass of amino acids concurrently expands, presumably as response to maintain fitness and protein function^10,11^. The strong N-limitation of phytoplankton primary production in the ocean is restricted to the permanently stratified mixed layer of tropical and subtropical regions^12^. In other regions and below the mixed layer different environmental and biotic drivers such as limitation by Carbon, other elements or temperature may control growth, the genomic G+C content and genome size of prokaryotic lineages^9,10,13^. However, there is no information available on the AGL, genomic G+C content, and N-ARSC of pelagic prokaryotic communities over ocean-wide latitudinal gradients with pronounced differences in N-availability and how they relate to nitrate concentrations, i.e. N-limitation of primary production.

## Results and Discussion

### Gene length of oceanic prokaryotes is a function of N availability

We assessed AGL, genomic G+C content and N-ARSC of the Atlantic Ocean Microbiome (AOM) over a 13,000 km transect from 62°S to 47°N covering regions of greatly varying N-availabilities (Fig. 1a-b, Supplementary Table 1). The transect included the tropics and subtropics where primary production is strongly N-limited (South Atlantic Gyre: SAG; North Atlantic Gyre: NAG), the temperate regions with seasonally fluctuating N-availabilities (South Atlantic: SA; North Atlantic: NA) and the Southern Ocean (SO) with permanently high nitrate concentrations^12,14^. Samples of the 0.2 to 3.0 μm-fraction collected at 22 stations at a depth of 20 m were paired-end shotgun Illumina sequenced resulting in a total of 206 Gb with a sample mean of 8.9+5.3 Gb (Supplementary Table 2). After assembly (total assembly length: >17.52 Gb), 12.05 million (M) gene sequences were predicted, and from these sequences we reconstructed the AOM reference gene catalogue (AOM-RGC) containing 7.75 M non-redundant (nr) protein-coding sequences, of which 56.6% were taxonomically classified. For the analysis of taxon resolved AGL, a subset of 3.67 M complete genes (55.7% taxonomically classified) were used. The majority of classified genes (83.9%) affiliated to Bacteria whereas minor proportions to Archaea, viruses and picoeukaryotes (16.1%). *Prochlorococcus, Synechococcus, Pelagibacteraceae, Rhodobacteraceae*, the SAR86 and SAR92 clades and *Flavobacteria* constituted the AOM to large extends, but supplemented with other lineages and exhibiting distinct biogeographic patterns (Supplementary Fig. 1). In total 38% percent of nr genes were functionally annotated by homology to a KEGG ortholog (KO). N-acquisition pathways comprised 0.85±0.19% of mapped reads at each station of which 54±6% encoded the glutamate synthase pathway. Amino acid transporters constituted 0.66±0.11% of all mapped reads. For a complete gene list see Supplementary Table 3.

**Figure 1.**
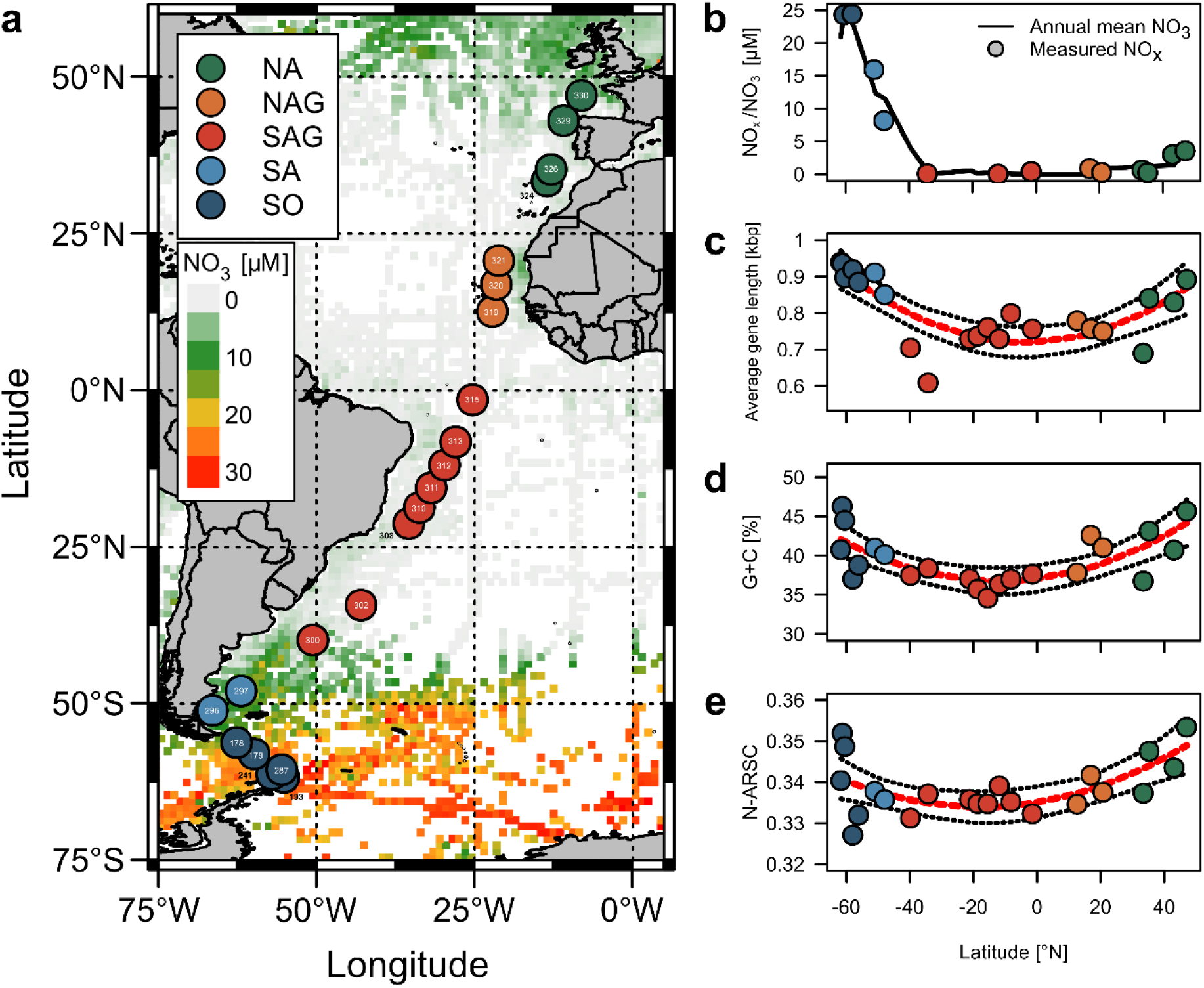
Stations and nitrate in the Atlantic and Southern Ocean visited during cruises ANTXXVIII/4 and -/5 with RV Polarstern and N-related genomic features of the Atlantic Ocean Microbiome (AOM). **a,** station location in the biogeographic regions SO, SA, SAG, NAG and NA (for abbreviations see text and for station details Table S1). Numbers of station are given in the circles and overlayed on a map with annual mean surface concentrations of nitrate (https://www.nodc.noaa.gov/OC5/woa18f/index.html). **b,** annual mean and ambient surface concentrations of nitrate and NO_x_ (nitrate+nitrite) in the biogeographic regions along the transect. **c-e**, distribution of AGL, genomic G+C content and N-ARSC in the biogeographic regions and their Spearman correlations (red line) and 95% confidence intervals (black dotted line) with latitude (AGL: r^2^: 0.58, p<0.001; G+C: r^2^: 0.46, p=0.001; N-ARSC: r^2^: 0.34, p=0.007).

The AGL exhibited a significant bimodal correlation with latitude with highest values in the SO and NA (Fig. 1c) and correlated also significantly with nitrate (Fig. 2e). A cluster analysis of normalized AGL of the 117 major prokaryotic genera and two virus families with ≥50 genes and ≥10 kb occurring at ≥50% of all stations showed two main AGL patterns over the transect (Supplementary Fig. 2). Cluster C1 (72% of tested taxa) encompassed all taxa with a bimodal correlation of AGL and latitude and exhibited a positive correlation with annual mean nitrate concentrations. AGL of taxa of this cluster were particularly small in the N-depleted (sub)tropics and included the Cyanobacteria *Prochlorococcus* and *Synechococcus*, the major alphaproteobacterial lineage *Pelagibacter* as well as the virus families *Podoviridae* and *Myoviridae* (Fig. 2a,b). AGL of this cluster exhibited significant correlations with latitude (r^2^=0.141, p<0.001) and nitrate (linear; r^2^=0.164, p<0.001). Taxa of cluster C2 (28% of tested taxa) exhibited no pronounced relationships with latitude (r^2^=0.04, p=0.001) or nitrate (r^2^=0.02, p<0.001) and encompassed genera like *Polaribacter* of Flavobacteria, *Planktomarina* of *Rhodobacteraceae* and the gammaproteobacterial SAR86 clade (Fig. 2c,d).

**Figure 2.**
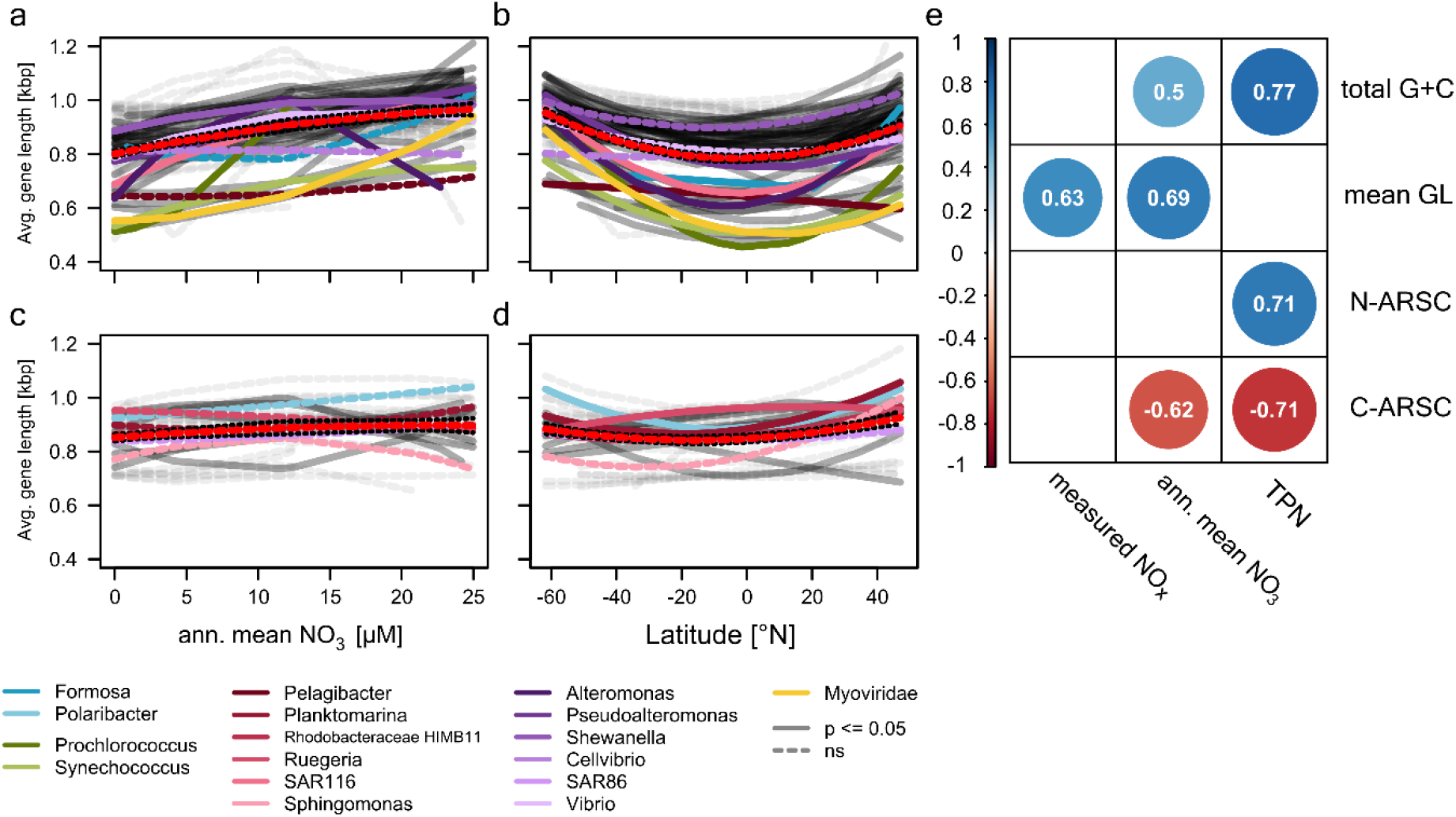
Correlation of AGL and genomic G+C content and N-ARSC of the AOM with latitude, nitrate and total particulate nitrogen (TPN). **a-d**, Correlations of AGL of clusters C1 and C2 encompassing major prokaryotic genera and virus families (see legend) with nitrate and latitude. Patterns were determined by using unimodal models. **e**, Correlation coefficients of Pearson correlations (p≤0.05) of AGL, genomic G+C content and N-ARSC with ambient and annual mean surface nitrate concentrations and TPN.

The number of prokaryotic genes and genome size over the entire size range of prokaryotic genomes has been shown to be positively correlated^2^. The prokaryotic AGL, however, has never been related to genome size, number of genes and genomic G+C content and it is unknown whether it varies with resource limitation. An analysis of these features of the 16,834 genomes available at NCBI (January 2020) yielded significant correlations between AGL, genome size and G+C content (Supplementary Figure 3). In oceanic systems, concentrations of inorganic nutrients and in particular of nitrate, often limiting phytoplankton primary production, can vary by orders of magnitude^12^. The positive correlation of AGL with nitrate concentration of many major genera of pelagic prokaryotes and two virus families of the AOM implies that N-availability or coupled growth constraints such as energy limitation drive the adaptive reduction in gene length with decreasing N-concentration. Hence, N-limitation appears to affect not only genome size, purifying selection towards a reduced genomic G+C content and N-ARSC of many marine bacterial genera^8–10^ but also AGL thus further lowering the N-demand and energy costs of biosynthetic reactions and DNA-replication. To compare the effect of saving N by reducing AGL, G+C content or N-ARSC we analyzed the theoretically reduced demand of N atoms required for nucleotides and amino acids as a function of transcription and translation cycles for these three variables in observed ranges occurring in marine prokaryotes (see above, Fig. 3). The outcome of this analysis demonstrates the dramatically higher effectiveness of reducing AGL than the genomic G+C content or N-ARSC in proteins for saving N. As transcription and translation cycles of individual genes may vary greatly and possibly irrespective of the growth phase it is difficult to exactly translate this effect of saving N to growth of individual prokaryotic lineages, but it demonstrates the great potential of this mode of saving N. This effect is particularly important at slow growth or during stationary phase at most severe resource limitations when maintenance metabolism predominates. Such conditions regularly occur in the most nutrient limited permanently stratified (sub)tropical oceanic gyres. Our results clearly show that reducing AGL is the most critical and not yet considered evolutionary mode of prokaryotes to adapt to N-limitation. Interestingly, it has been shown that an abundant marine prokaryote responds to N limitation also on the transcriptional level. Under N-deplete conditions *Prochlorococcus* starts transcribing various genes downstream of the transcriptional start site more frequently than under N-replete conditions leading to a reduced demand of N and other resource in the transcripts^15^. Such a reduction of the transcribed gene regions may lead to reducing the gene length to a size ensuring the functionality of the encoded protein as evolutionary adaptation to N-deprived conditions.

**Figure 3.**
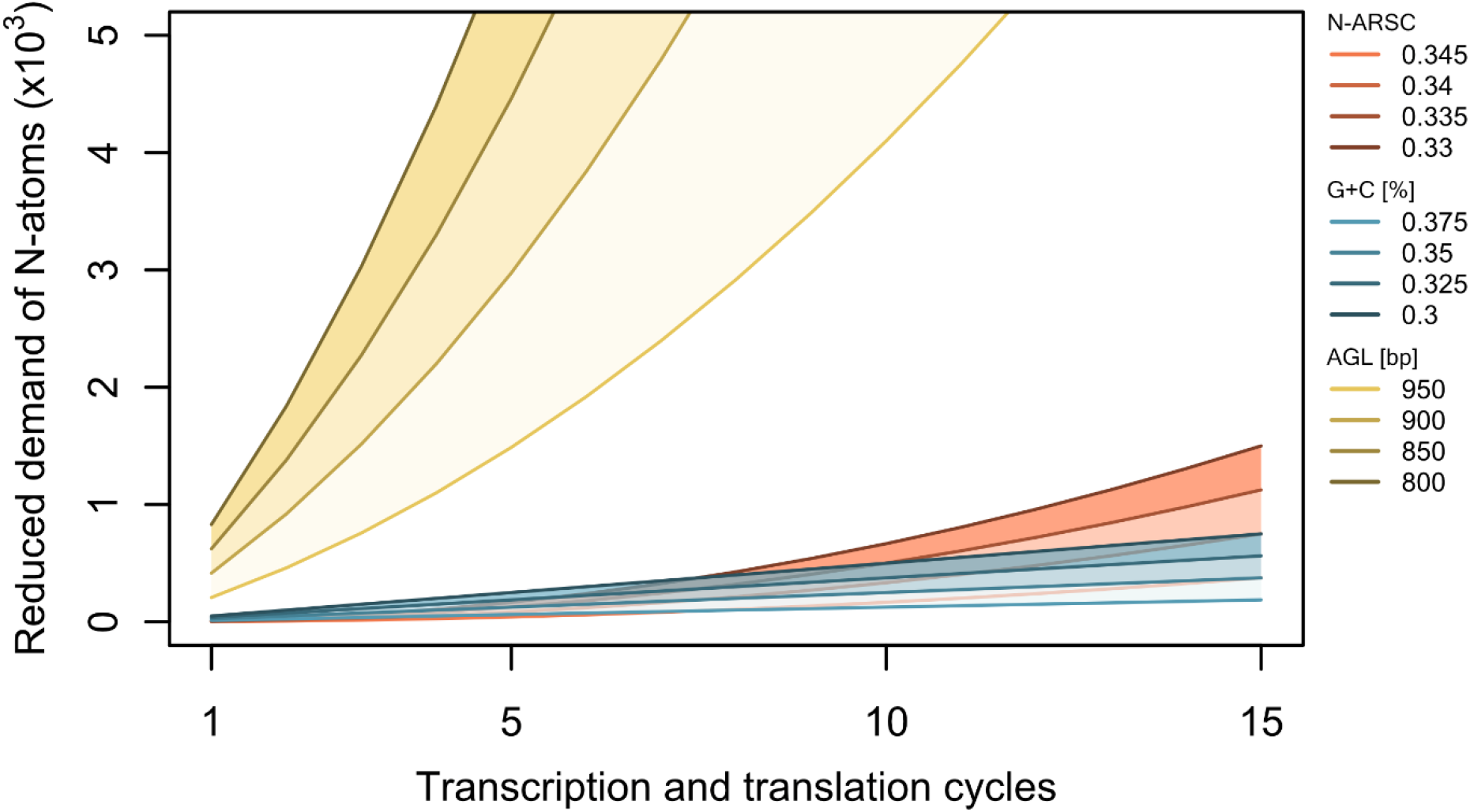
Reduction of N demand for given ranges of gene length, genomic G+C content and N-ARSC over increasing numbers of transcription and translation cycles. Numbers of N atoms saved were calculated for the given values of N-ARSC, G+C content and gene length for nucleotides needed for the transcription and translation cycles and amino acids produced for protein synthesis based on a reference gene length of 1000 bp, G+C content of 40% and an N-ARSC of 0.35. These reference values were in the range of values occurring in N-replete regions of our data set.

As AGL of the genera of the *Pelagibacteraceae* (Fig. 2a,b) is particularly small this indicates that genome streamlining of this prominent oceanic family does not only lead to a reduction in gene number^2,4^ but also in gene length. Our findings have most interesting evolutionary implications. Mutation, purifying selection, HGT and gene loss are well known mechanisms of the adaptive evolution of genes and genome streamlining enhancing metabolic efficiency of the evolving populations at the prevailing environmental conditions^4^. Reduction of gene length, does not only occurring in prokaryotes but also in bacteriophages, presumably in their state as prophage, is a novel mechanism of genome reduction but its mode of action is unknown. We speculate that it may act by non-synonymous or synonymous mutation and subsequent codon deletion, deletion of a gene fragment downstream of the transcriptional start site or replacement of genes by homologs of reduced size via HGT while maintaining the metabolic efficiency of the encoded protein at an optimal or sub-optimal but acceptable level as a trade-off. An important follow-up question is to examine whether different genes underlie similar constraints of reducing their size and whether this is taxon-specific or a general feature. Future work is needed to mechanistically understand the molecular basis of evolutionary reduction of gene length in prokaryotes and phages under environmental constraints of N-availabilities and possibly energy limitation.

### G+C content and N-ARSC of the AOM

The G+C content of all genes and the proteomic N-ARSC exhibited also bimodal latitudinal patterns (Fig. 1d,e). Lowest values consistently occurred in the permanently stratified SAG and the highest values in the SO and NA. The G+C content correlated positively with annual mean nitrate and N-ARSC with measured nitrate concentrations and total particulate N (Fig. 2e). G+C content and N-ARSC correlated also positively (Pearson correlation 0.84, p <0.001), in line with a global trend including all prokaryotic genomes (Supplementary Fig. 3).

The analysis of the normalized genetic G+C content of marine prokaryotes and several virus families over the transect yielded four distinct patterns grouped into clusters (Fig. 4a-d). Clusters C1 (41.1 % of all taxa) and C2 (19.3%) showed a general increase of G+C with ambient nitrate concentration. Both showed highest G+C values in the SO and SA and a decrease in the (sub)tropics. In contrast to cluster C2, C1 exhibited a minor G+C increase in the NA. Both clusters encompassed mainly Alpha- and Gammaproteobacteria (Fig. 4i) but also other major genera such as *Prochlorococcus*, indicating a subclass-specific adaptation strategy. Cluster C3 (16.4%) showed an inverse G+C distribution with high values in the N-depleted SAG and NAG and low values in the SO; it included *Synechococcus* and the SAR116 clade and other genera of generally low or distinct regional abundance (Fig. 4e,f, Supplementary Fig. 1). G+C of genera belonging to cluster C4 (23.2%) did not show a consistent relationship with ambient nitrate concentration but exhibited two distinct peaks in the SA/SAG and NAG (Fig. 4g,h). C4 consisted mainly of Flavobacteriia (Fig. 4i) as well as genome-streamlined genera of *Pelagibacteraceae* with an overall low G+C content. These data indicate that lineages known to be active players of prokaryotic communities in various oceanic regions^16–21^ have a relatively low G+C content in SAG and NAG. All of them, however, exhibit the highest G+C content in SO where growth is usually not N-limited. These patterns, exhibited by the majority of taxa, are consistent with the concept of an adaptive evolution towards a reduced G+C content under N-limiting conditions^9,13^ even though the mechanisms involved remain unclear. Mutation bias towards a reduced G+C content and purifying selection but other not fully understood selective forces^3,8^ seem to be involved. The fact that the selective forces act preferentially on actively transcribed protein coding genes^6^ may explain why predominantly the more prominent and active players of the prokaryotic communities exhibit these patterns. The lineages with a permanently low G+C content underlie other selective processes, genome streamlining as they exhibit the smallest genomes^2^ and AGL (see above). However, as the genome size is positively correlated with the G+C content in the larger phylogenetic groups to which the major lineages of marine pelagic prokaryotes affiliate^7^ and also when considering all available genomes (Supplementary Fig. 3), genome streamlining appears to be inherently associated with reducing the G+C content. The other lineages with no reduced G+C content in the most strongly N-limited SAG presumably underlie other evolutionary constraints than N-limitation. They are either minor components of the microbial communities with generally little activity and thus a presumably low adaptive evolutionary forcing towards a reduced G+C content^6^. Or they underlie other forces when they dwell predominantly in less N-depleted regions than SAG, such as *Synechococcus*, or occupy niches with no N-limitation, such as Planktomycetes and Verrucomicrobia, on N-rich particles and Carbon limitation^5,13,22,23^.

**Figure 4.**
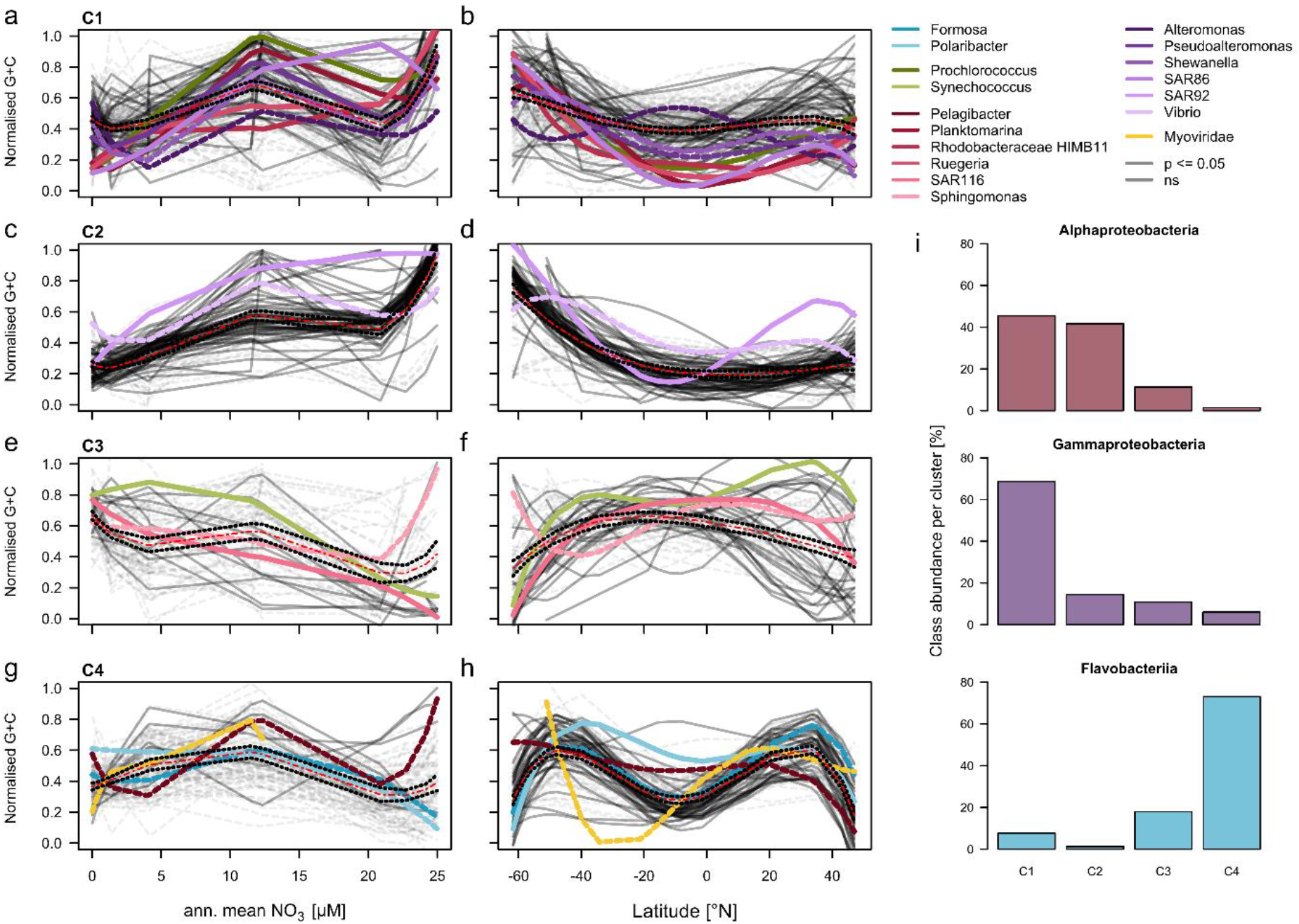
Patterns of the genomic G+C content of the AOM. **a-h,** Correlations of the normalized G+C content of four clusters (C1, C2, C3, C4) of major prokaryotic genera and virus families (see legend) in the Atlantic and Southern Ocean in correlation to annual surface nitrate concentration and latitude. Patterns were determined by using non-linear models. Fit significance of each genus is indicated by a solid line (for further details see legend). **i**, Affiliation of major classes of prokaryotes to clusters C1 to C4 of the G+C correlation patterns.

### Diversity and patterns of genes involved in N-acquisition

As availability of N is crucial for oceanic prokaryotes and the evolution of their genomic structure (see above) an important and related question is in which forms N is available to prokaryotes under various nutrient regimes. Besides oxidized inorganic forms and neutral gas a variety of reduced N containing compounds exist (e.g. oligopeptides, amino acids, urea). These are the preferred N-sources of heterotrophic prokaryotes because reduction of the oxidized forms is energy-costly (Fig. 5a). Break down products of proteins originating predominantly from phytoplankton such as oligopeptides, dipeptides and free amino acids are major N-sources of heterotrophic and partly of autotrophic pelagic prokaryotes (Fig. 6a). Ammonium and urea act as important metabolites of the amino acid and pyrimidine metabolism and can be important N-sources^24–28^. Other reduced organic N-species including cyanate may be available as well^29^ (Fig. 5a). There is some information on the genetic potential and proteomic spectrum of pelagic prokaryotes to acquire inorganic and organic N-species in pelagic ecosystems^9,17,18,29,30^. However, we still lack a comprehensive and detailed insight into the genetic potential of pelagic prokaryotic communities to acquire potentially available inorganic and organic N-species on a global or ocean-basin scale including regions with and without strong N-limitation of phytoplankton primary production. This is particularly important considering the utmost relevance of N in shaping genomic traits of prokaryotes in oceanic systems under N-limitation (see above) and strong competition for and exploitation of available N-sources. Therefore, we screened the AOM for key genes involved in the acquisition of the entire range of N-species. The AOM harbors a large variety of genes encoding transporters of oligo- and dipeptides, various amino acids, alkylamines, ammonium, cyanate, formamide, nitriles and the metabolism of urea and oxidized N-species (Fig. 6a,b). Richness and effective number of these genes was lowest in the N-replete SO (Supplementary Fig. 6). The other N-depleted regions exhibited some variations but no distinct patterns. Although many pathways showed only very low abundances, biogeographic patterns as well as phylogenetic affiliations were visible. Glutamate synthase, leading to the final step of intracellular ammonium transfer for further metabolic reactions, comprised 54±6% of the N-acquisition genes and was highly abundant in the stratified SAG and NAG. Transporters of branched chain and general amino acids, glycine betaine/proline, octopine/nopaline, oligopeptides, ammonium, urea and urease and glutamate synthase constituted the great majority of these genes (Fig. 6a). Genes encoding transporters of general amino acids, urea and ammonium and urease exhibited highest abundances in SAG whereas those encoding transport of oligopeptides were most abundant in SO. Genes encoding transporters of glycine betaine/proline, branched chain amino acids and glutamate synthase showed a more patchy or rather even distribution over the transect (Fig. 6a). Among the less abundant genes cyanate lyase and nitrilase showed highest abundances in the SO (Fig. 5b). In general, Alphaproteobacteria exhibited the largest variety of N-acquisition genes, followed by Gammaproteobacteria (Fig. 6b). Several prokaryotic classes dominated or were distinct for specific N-acquisition genes: Gammaproteobacteria for transporters of oligopeptides and several individual amino acids, Betaproteobacteria for denitrification and transporters of glutamate/aspartate, Bacteroidetes/Flavobacteria for nitrilase, *Synechococcaceae* for assimilatory nitrate and nitrite reduction and *Prochloraceae* for urea transporters and urease (Fig. 6b, Supplementary Fig. 7). Hence, the latitudinal distribution of these phylogenetic groups was closely linked to that of the respective genes. Distribution of genes encoding amino acid transporters among Alphaproteobacteria reflected the relative abundances of the various families (Supplementary Fig. 8). However, genes encoding transporters of dipeptides, ammonium transporters and urease were specifically dominated by distinct families with variation in the different regions (Supplementary Figs. 7, 8).

**Figure 5.**
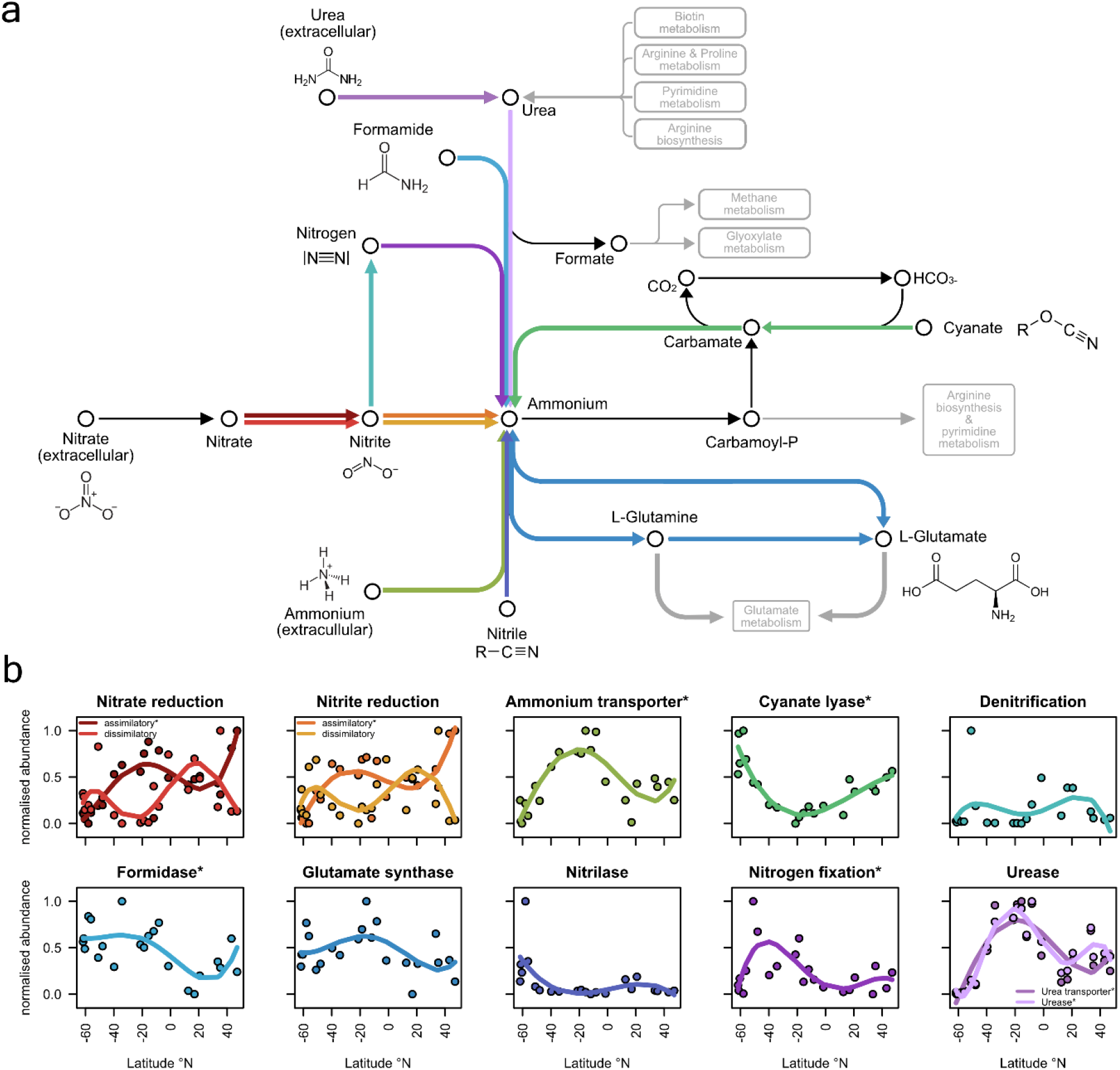
N-acquisition genes and their distribution along the Atlantic Ocean transect. **a,** pathways of N-acquisition genes leading to intracellular ammonium. **b**, normalised distribution of N-acquisition genes along the transect.

**Figure 6.**
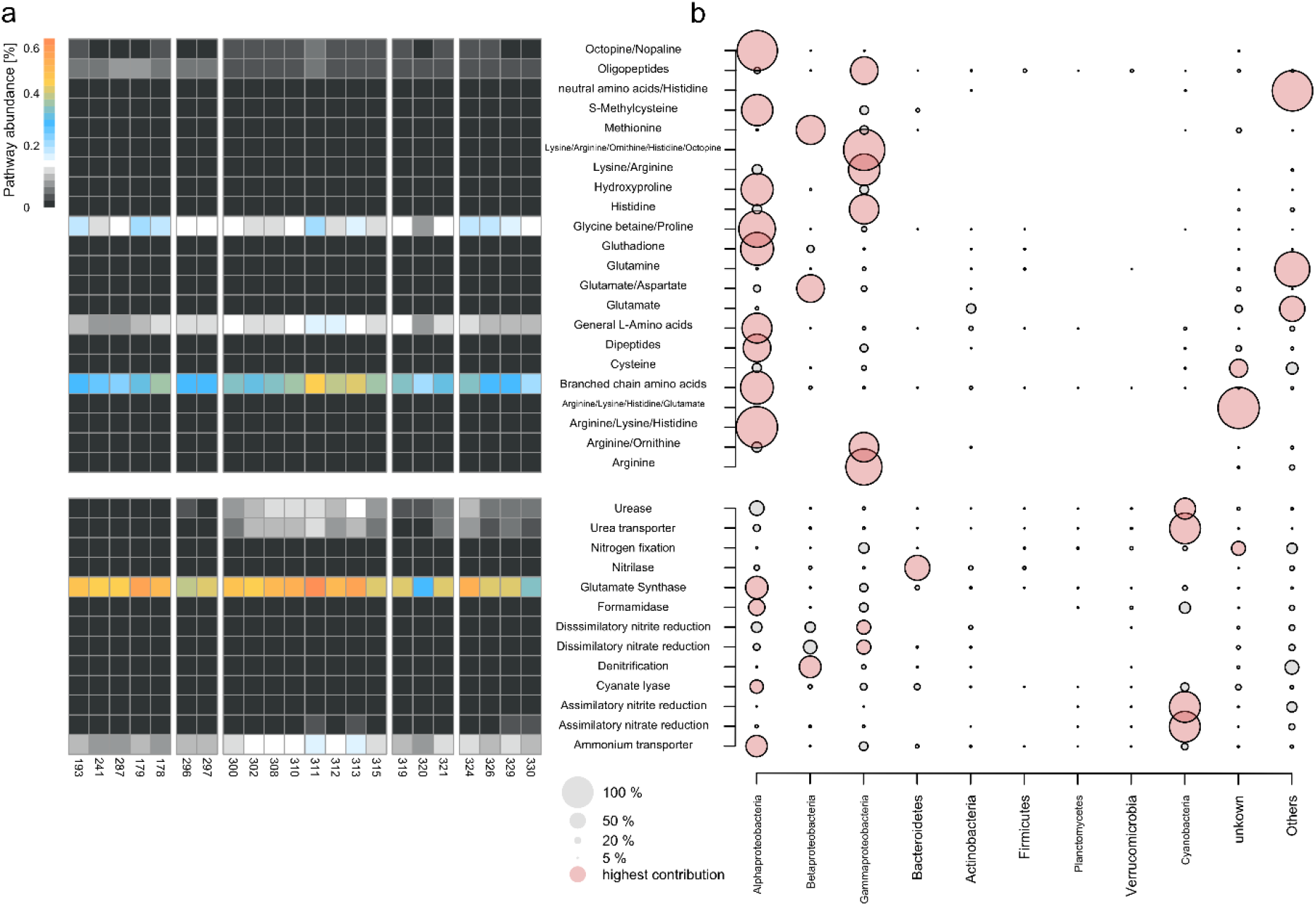
N-acquisition genes along the transect and their distribution among the prokaryotic groups of the AOM. **a,** relative abundance of genes encoding transporters of amino acid-related and other organic N-compounds at stations along the transect. **b**, relative distribution pathways and transporters among prokaryotic phyla and classes. For higher phylogenetic resolution of major lineages see supplementary Fig. S7, S8.

The results demonstrate that the AOM’s repertoire of genes acquiring N-containing compounds is very diverse thus enabling its different members to occupy many niches to exploit the large spectrum of potentially available N-species. The highest diversity of these genes existed in regions with strong N-limitation, presumably driving this diversification because of the high competition for this very precious element The different patterns of N-acquisition genes in the various prokaryotic families indicate that overlap of different families to exploit N-containing compounds was rather limited thus emphasizing different strategies and niches for the acquisition of N among free-living bacteria, presumably reducing functional redundancy. This notion is in line with a recent concept of reduced functional redundancy due to these auxiliary genetic features not considered in KO categories^31^. Whereas most N-acquisition genes and their affiliation to phylogenetic groups were expected based on previous findings^17,18,29^ it was unexpected to find genes encoding cyanate lyase affiliated to quite different bacterial phylogenetic groups. Concentrations of cyanate in pelagic systems are in the nM range and incorporation can meet up to 10% of total N-demand^32^. So far, use of cyanate has been attributed mainly to ammonium-oxidizing prokaryotes^29^ but our finding of genes encoding cyanate lyase along the transect suggests that cyanate is used as an N-source for biosynthetic requirements by other phylogenetic lineages but needs to be tested. It was also surprising to find optopine/nopaline transporter-like sequences, especially in *Pelagibacteraceae*. Both are derivatives of the amino acids glutamate, arginine and alanine and their transport systems are known from *Agrobacterium tumefaciens*. Both compounds are produced by the host plants after infection by the virulence plasmid to promote growth of tumors and the production of secondary metabolites^33^. Genes encoding these transporters have not been described in marine prokaryotes. Whether our finding is based on incorrect gene annotation or indicates that these transporters may also mediate uptake of other similar compounds needs to be tested.

### Conclusion

Our investigation shows that availability of N in the form of nitrate and related biogeochemical effects such as limitation of phytoplankton primary production have fundamental effects on shaping genomic and proteomic traits of the microbiome of the sunlit Atlantic Ocean. Shifts in community wide AGL and G+C content were not caused by a changing community structure but were evident for many and in particular major phylogenetic groups which exhibited respective variation in the genomic G+C content, N-ARSC and as a completely novel finding AGL. AGL has the greatest effect on reducing the N-demand and can vary by approximately 20% within a narrow phylogenetic range. In response to the sparse availability of N in particular in the N-depleted regions a highly diverse repertoire of N-acquisition genes in the different prokaryotic families is present which enables the AOM to maximize N-acquisition. The discovery that AGL is affected by N-availability presumably is not restricted to oceanic prokaryotic communities but may be a consequence of N-deficiency also in other N-limited prokaryotic communities.

Methods, along with any additional Extended Data display items and Source Data and related references, are available in the online version of the paper.

## Acknowledgements

We thank the master, his crew and the principal scientists (M. Lucassen, K. Bumke) of cruises ANT XXVIII/4 and -/5 of RV Polarstern, T.H. Badewien, A. Gavrilov, S. Rackebrandt, T. Remke, J. Vollmers, M. Wietz, I. Wagner-Döbler and M. Wurst for cruise support, M. Heinemann, B. Kuerzel, R. Weinert and C. Lehners for technical laboratory assistance. Constructive suggestions by S. Biller on an earlier version of this manuscript are gratefully acknowledged. This work was funded by Deutsche Forschungsgemeinschaft within the Collaborative Research Center *Roseobacter* (TRR51) and the Graduate Research training group “The Ecology of Molecules” (EcoMol) supported by the Lower Saxony Ministry for Science and Culture.

## Author contribution

LD carried out the bioinformatics and statistical analyses and wrote the draft of the publication; AP and BP carried out the metagenomics sequencing and quality control of the raw sequences; BW carried out sampling and sample filtration; HAG analyzed nitrate concentrations; RD supervised the metagenomics sequencing and contributed to reviewing the manuscript; MS designed the study, supervised the bioinformatics and statistical analyses and finalized the draft manuscript. All authors reviewed the manuscript.

## Methods

Twenty-two stations between 62°S and 47°N were visited during cruises ANT XXVIII/4, 13 March–9 April 2012, and ANT XXVIII/5, 10 April–15 May 2012, with RV Polarstern. For exact locations of the stations see Table S1. Samples were collected at 20 m depth with 12 l-Niskin bottles mounted on a Sea-Bird Electronics SBE 32 Carousel Water Sampler equipped with a temperature, salinity, depth probe (SBE 911 plus probe), a chlorophyll fluorometer (Wet Labs ECO – AFL/FL) and transmissometer (Wet Labs C-Star). Nitrate concentration was analyzed in prefiltered (0.2 μm, isopore membrane filter, EMD Millipore Corporation, USA) and HgCl2-preserved and frozen (−20°C) subsamples in the home lab after thawing using a microtiter plate reader (FLUOstar Optima, BMG Labtech, Germany) following established procedures for N oxides (NOx)^34^. For the analysis of total particulate N (TPN), 1– 4 l of seawater were filtered through Whatman GF/F filters and stored at −20°C until analysis in the home lab. POC and TPN were analyzed as described previously^35^. For metagenomics analysis, water of several bottles was pooled in an ethanol-rinsed polyethylene barrel to a total volume of 40 l. Within 60 min after collection the sample was prefiltered through a 10-μm nylon net and a filter sandwich consisting of a glass fiber filter (47 mm diameter, Whatman GF/D; Whatman, Maidstone, UK) and 3.0-μm polycarbonate filter (47 mm diameter, Nuclepore; Whatman). Picoplankton was harvested on a filter sandwich consisting of a glass fiber filter (47 mm diameter, Whatman GF/F; Whatman) and 0.2-μm polycarbonate filter (47 mm diameter, Nuclepore; Whatman). All filters were immediately frozen in liquid N and stored at −80°C until further processing. Environmental DNA was extracted from the filter sandwich and subsequently purified employing the peqGOLD gel extraction kit (Peqlab, Erlangen, Germany) as described previously^36^. Illumina shotgun libraries were prepared using the Nextera DNA Sample Preparation kit as recommended by the manufacturer (Illumina, San Diego, USA). To assess quality and size of the libraries, samples were run on an Agilent Bioanalyzer 2100 using an Agilent High Sensitivity DNA kit as recommended by the manufacturer (Agilent Technologies, Waldbronn, Germany). Concentrations of the libraries were determined using the Qubit^®^ dsDNA HS Assay Kit as recommended by the manufacturer (Life Technologies GmbH, Darmstadt, Germany). Sequencing was performed by using the HiSeq2500 instrument (Illumina Inc., San Diego, USA) using the HiSeq Rapid PE Cluster Kit v2 for cluster generation and the HiSeq Raid SBS Kit (500 cycles) for sequencing in the paired-end mode and running 2×250 cycles.

### Annual mean nitrate concentrations

Annual mean nitrate concentrations at 20 m depth of each station were extracted from the 1° World Ocean Atlas 2009, provided by the National Oceanic and Atmospheric Administration (https://www.nodc.noaa.gov/cgi-bin/OC5/woa18f/).

### Metagenomic assembly and gene prediction

Illumina reads were quality checked and low-quality regions as well as adaptor sequences were trimmed using Trimmomatic 0.36^37^ (*ADAPTER:2:30:10 SLIDINGWINDOW:4:25 MINLEN:100*). The high quality (HQ) reads were assembled using metaSPAdes 3.11.1^38^. Contigs smaller than 210 bp and average coverage <2 were discarded. Gene-coding sequences of the assembled contigs were predicted using Prodigal 2.6.2 in meta-mode^39^. Genes shorter than 210 bp and longer than 4,500 bp were discarded to account for prokaryotic and eukaryotic gene length. This resulted in 8.38 M partial and 3.67 M complete unique gene sequences (supplementary table S2).

### Taxonomic classification of gene sequences

Gene sequences were taxonomically classified using Kaiju 1.6^40^ (-*greedy* mode with 5 allowed substitutions and e-value 10e-5) and the NCBI nr database (downloaded on 2018-05-29) including prokaryotic, eukaryotic and viral sequences as well as the proGenomes database^41^ (downloaded on 2019-07-26). Gene taxonomy was compared between both approaches and last known ancestor was inferred from the highest available phylogenetic resolution. In total 63.4% of all unique genes and 55.7% of complete genes were taxonomically classified.

### Gene catalogue generation

To generate a non-redundant (nr) gene catalogue, gene sequences were clustered at 95% identity using USEARCH 10.0.24^42^ (*-cluster_fast–id 0.95*). The resulting 7.75 M cluster centroids were used as representative AOM gene sequences. Genes were taxonomically classified as described above. Gene functions were assigned using the Kyoto Encyclopedia of Genes and Genomes (KEGG) online annotation tool GhostKOALA^43^ (https://www.kegg.jp/ghostkoala/) using the prokaryotic, eukaryotic and viral KEGG gene database (release 86) and default settings. In total, 59% of genes were taxonomically classified and 39% of all sequences were assigned to a KEGG orthologue (KO).

### Illumina read abundance and normalization

To acquire gene abundance data, HQ Illumina reads longer than 75 bp were mapped to the AOM gene sequences using bowtie2^44^ 2.3.5 (--*very-sensitive-local* mode). SAMtools^45^ version 1.9-58-gbd1a409 was used to convert the SAM alignment file to read abundance tables. Reads that did not map to any nr sequence were discarded. To account for different sequencing depth and gene length, counts from each station were normalized by dividing read counts by gene length in kb to obtain reads per kilobase (RPK). Subsequently scaling factors were calculated for each sample by dividing the sum of RPKs by one million. The scaling factors were used to normalize the RPKs of each sample to counts per million (CPM)^46^

### Determination of G+C, N/C-ARSC, molecular protein weight and gene length

To determine G+C content, Nitrogen/Carbon content of amino acid residual side chains (N/C-ARSC) genes predicted from individual stations were classified as described above. G+C content of each predicted gene was determined by dividing the total amount of G and C bases by the total gene length. To determine N- and C-ARSC, nucleotide sequences were translated to amino acids. Average N and C content of amino acid side chains was calculated for every gene according to sum formula of each amino acid. The same approach of gene prediction and determination of genomic traits was applied to 16,834 complete bacterial reference genomes downloaded from NCBI GenBank (January 2020).

### Statistical analysis

All statistical evaluations were performed in R (version 3.6.0; https://www.r-project.org/) using the additional packages *vegan*^47^ *(v2.5-6), ape*^48^ *(v5.3)*, and *cluster*^49^ *(v2.0.8)*.

### AGL, G+C content, and N-ARSC

Patterns of AGL were analyzed on the level of complete genes of prokaryotic genera and virus families. For a similar analysis of the G+C content also incomplete genes were included. Only taxa with more 50 genes (min. 10kb) in ≥50% of all samples were considered. Stations with less than 50 genes were excluded in the analysis of each taxon.

To compare trends across genera, AGL and G+C content were normalized to values between 0 and 1 (formula: x-min(x)/max(x)-min(x)). Euclidean distances of G+C and AGL profiles were calculated and subsequently clustered using minimal variance Ward.D2 clustering. Linear/non-linear model fitting was used to determine a relationship between G+C and AGL to annual mean nitrate concentration for each resulting cluster. Correlations between AGL, G+C, and N/C-ARSC and environmental parameters were calculated from all stations where data with environmental data were available (Table S1). P-values ≤0.05 were considered significant.

To analyse geographic distribution patterns, abundances of genes involved in N-acquisition (Supplementary Table 3) were normalised to values between 0 and 1 (see above). Effective number of the same genes was calculated after Jost 2006^50^.

### Data availability

Sequence data were deposited under the INSDC accession number PRJEB34453 in the European Nucleotide Archive (ENA) using the data brokerage service of the German Federation for Biological Data^51^ (GFBio), in compliance with the Minimal Information about any (X) Sequence (MIxS)^52^ standard. Environmental data of the stations and depth collected during cruises ANTXXVIII/4 and −/5 are available at https://doi.pangaea.de/10.1594/PANGAEA.906247.

## Supplementary Material

### Tables

**Table S1** Station details, hydrography, nitrate, G+C, AGL and N/C-ARSC-data

**Table S2** Sequencing and assembly statistics of the Atlantic Ocean Metagenomes

**Table S3** List of genes encoding proteins of N-acquisition and AA-transport

### Figures

**Figure S1:** Heatmap of Relative abundances of prominent taxa in the Southern and Atlantic Ocean

**Figure S2:** Heatmap of taxonomically resolved AGL data and clusters

**Figure S3:** Correlation of genome size and G+C content, genome size and G+C content (NCBI-data of available genomes).

**Figure S4**: Dendrogram based on patterns of G+C distribution among prokaryotic genera and virus families

**Figure S5**: Heatmap of taxonomically resolved G+C data

**Figure S6**: Richness and EN of N-acquisition genes

**Figure S7**: Taxonomically resolved abundances of N-acquisition pathways

**Figure S8**: Taxonomically resolved abundances of amino acid transport systems

