## Supplementary Material - 3 Tables+8 Figures for "Nitrogen availability drives gene length of dominant prokaryotes and diversity of genes acquiring Nitrogen-species in oceanic systems"

**Supplementary Information**

**Table S1** Station details, hydrography, nitrate, G+C, AGL and N/C-ARSC-data

**Table S2** Sequencing and assembly statistics of the Atlantic Ocean Metagenomes

**Table S3** List of genes encoding proteins of N-acquisition and AA-transport

**Supplementary Figures**

**Figure S1:** Heatmap of Relative abundances of prominent taxa in the Southern and Atlantic Ocean

**Figure S2:** Heatmap of taxonomically resolved AGL data and clusters

**Figure S3**: Correlation of genome size and G+C content, genome size and G+C content (NCBI-data of available genomes).

**Figure S4**: Dendrogram based on patterns of G+C distribution among prokaryotic genera and virus families

**Figure S5**: Heatmap of taxonomically resolved G+C data

**Figure S6**: Richness and EN of N-acquisition genes

**Figure S7**: Taxonomically resolved abundances of N-acquisition pathways

**Figure S8**: Taxonomically resolved abundances of amino acid transport systems

**Table S1**

Station number, latitude, longitude, date of visit, temperature, salinity, chlorophyll *a* (Chl *a*), total particulate Nitrogen (TPN), measured NOx (nitrate+nitrite) and annual mean nitrate concentrations at the depth of sampling (20 m) during cruises ANTXXVIII/4 and -/5 with RV Polarstern.

| Station | Date [m/d/y] | Longitude [°W] | Latitude [°N] | Temperature [°C] | Salinity [psu] | Chl a [µg L-1] | TPN [µg L-1] | $NO_x$ [µM] | $NO_3^-$ ann. mean [µM] | avg. G+C [%] | AGL [bp] | N-ARSC | C-ARSC |
|---|---|---|---|---|---|---|---|---|---|---|---|---|---|
| **193** | 03/19/2012 | -55.167 | -61.717 | 1.225 | 34.195 | 0.800 | 0.025 | *NA* | 20.881 | 40.745 | 939.487 | 0.340 | 2.906 |
| **241** | 03/25/2012 | -57074 | -61.244 | 1.226 | 34.043 | 1.186 | 0.026 | *NA* | 24.981 | 46.275 | 935.313 | 0.352 | 2.840 |
| **287** | 04/04/2012 | -55.500 | -60.500 | 1.832 | 33.837 | 1.229 | 0.030 | 24.300 | 24.151 | 44.484 | 897.071 | 0.349 | 2.867 |
| **179** | 03/16/2012 | -59.867 | -57.983 | 4.392 | 33.707 | 0.078 | 0.007 | 24.400 | 22.700 | 37.015 | 919.511 | 0.327 | 2.940 |
| **178** | 03/16/2012 | -62.633 | -56.170 | 6.981 | 33.979 | 0.347 | 0.012 | *NA* | *NA* | 38.766 | 884.332 | 0.332 | 2.918 |
| **296** | 04/11/2012 | -66.459 | -51.054 | 8.148 | 33.001 | 0.397 | 0.014 | 15.940 | 12.337 | 40.976 | 910.624 | 0.338 | 2.911 |
| **297** | 04/12/2012 | -61.923 | -47.941 | 10.344 | 33.492 | 0.565 | 0.022 | 8.170 | 11.485 | 40.146 | 850.815 | 0.336 | 2.899 |
| **300** | 04/15/2012 | -50.577 | -39.798 | 19.484 | 35.311 | 0.297 | 0.011 | *NA* | 3.998 | 37.467 | 704.164 | 0.331 | 2.925 |
| **302** | 04/17/2012 | -42.965 | -34.239 | 22.131 | 35.843 | 0.145 | 0.006 | 0.080 | 0.041 | 38.393 | 608.961 | 0.337 | 2.937 |
| **308** | 04/21/2012 | -35.402 | -21.229 | 27.149 | 37.316 | 0.098 | 0.006 | *NA* | 0.568 | 37.042 | 729.280 | 0.336 | 2.951 |
| **310** | 04/22/2012 | -33.725 | -18.664 | 27.371 | 37.394 | 0.106 | 0.006 | *NA* | 0.130 | 35.713 | 736.882 | 0.335 | 2.967 |
| **311** | 04/23/2012 | -31.862 | -15.522 | 26.882 | 37.240 | *NA* | *NA* | *NA* | 0.253 | 34.597 | 760.364 | 0.335 | 2.991 |
| **312** | 04/24/2012 | -29.752 | -11.898 | 27.270 | 37.014 | 0.057 | 0.006 | 0.040 | 0.307 | 36.329 | 729.655 | 0.339 | 2.979 |
| **313** | 04/25/2012 | -27.991 | -8.211 | 28.012 | 36.285 | 0.087 | 0.006 | NA | 0.051 | 36.998 | 799.001 | 0.335 | 2.960 |
| **315** | 04/27/2012 | -25.318 | -1.487 | 27.826 | 35.977 | 0.330 | 0.011 | 0.410 | 0.034 | 37.641 | 756.230 | 0.332 | 2.928 |
| **319** | 05/01/2012 | -22.196 | 12.590 | 22.923 | 35.776 | 0.722 | *NA* | *NA* | *NA* | 37.801 | 778.945 | 0.335 | 2.944 |
| **320** | 05/02/2012 | -21.569 | 16.904 | 21.415 | 36.206 | 0.506 | 0.012 | 0.890 | 0.000 | 42.594 | 756.844 | 0.342 | 2.896 |
| **321** | 05/03/2012 | -21.170 | 20.705 | 20.770 | 36.815 | 0.450 | 0.014 | 0.280 | 0.876 | 41.013 | 749.458 | 0.338 | 2.911 |
| **324** | 05/07/2012 | -13.534 | 33.398 | 17.787 | 36.606 | 0.146 | 0.009 | 0.660 | *NA* | 36.741 | 689.743 | 0.337 | 2.970 |
| **326** | 05/08/2012 | -12.897 | 35.239 | 17.918 | 36.619 | 0.214 | 0.009 | 0.240 | *NA* | 43.129 | 840.833 | 0.348 | 2.915 |
| **329** | 05/10/2012 | -10.935 | 43.042 | 13.744 | 35.786 | 0.735 | 0.019 | 2.990 | 1.390 | 40.702 | 829.834 | 0.344 | 2.924 |
| **330** | 05/11/2012 | -8.091 | 47.044 | 12.579 | 35.649 | 1.902 | 0.052 | 3.560 | 4.182 | 45.670 | 892.263 | 0.353 | 2.884 |

**Table S2**

Statistics of sequencing, assembly and ORF-prediction of the AOM samples of all stations visited and analyzed. *: based on contigs ≥ 210 bp contigs and ORFs, **: based on contigs ≥ 500 bp

| Station | Total bp | Joined reads | Contigs* | Contigs** | N50** | Total assembly length [bp]* | Largest contig [bp] | Predicted ORFs | Complete genes |
|---|---|---|---|---|---|---|---|---|---|
| **193** | 7,435,000,546 | 12,427,245 | 1,501,196 | 183,054 | 1,654 | 606,362,663 | 115,786 | 417,100 | 115,631 |
| **241** | 6,071,598,618 | 10,027,320 | 962,824 | 116,387 | 1,571 | 387,239,824 | 325,929 | 268,873 | 85,742 |
| **287** | 6,322,636,050 | 9,471,523 | 1,434,266 | 183,574 | 1,126 | 555,042,209 | 258,892 | 333,526 | 105,100 |
| **179** | 6,198,682,756 | 7,599,385 | 971,371 | 143,473 | 1,874 | 436,536,877 | 262,003 | 290,543 | 122,218 |
| **178** | 6,064,805,297 | 8,176,910 | 1,300,260 | 130,039 | 1,509 | 486,212,585 | 306,237 | 287,516 | 93,055 |
| **296** | 19,592,095,808 | 26,002,512 | 2,505,074 | 427,057 | 1,894 | 1,203,911,331 | 256,578 | 100,1702 | 334,728 |
| **297** | 6,616,159,289 | 9,041,391 | 1,357,387 | 198,801 | 1,617 | 585,795,325 | 162,636 | 441,554 | 132,975 |
| **300** | 6,792,243,498 | 12,739,153 | 1,582,103 | 360,444 | 877 | 780,657,750 | 143,583 | 567,271 | 118,897 |
| **302** | 5,503,346,589 | 11,099,908 | 1,398,405 | 322,151 | 807 | 675,992,337 | 81,215 | 258,162 | 104,395 |
| **308** | 4,841,364,911 | 8,620,757 | 1,087,405 | 263,756 | 1,044 | 578,609,656 | 152,476 | 388,272 | 130,983 |
| **310** | 6,517,496,619 | 11,188,810 | 1,220,934 | 292,443 | 1,033 | 637,971,540 | 93,510 | 494,727 | 134,333 |
| **311** | 7,559,399,537 | 11,548,284 | 1,276,149 | 323,727 | 1,029 | 676,526,273 | 154,832 | 549,470 | 136,285 |
| **312** | 4,447,667,609 | 7,750,178 | 1,055,905 | 241,878 | 1,040 | 550,785,256 | 125,412 | 368,056 | 115,755 |
| **313** | 7,892,259,228 | 13,906,649 | 1,416,050 | 310,566 | 1,068 | 729,545,280 | 782,123 | 548,478 | 142,712 |
| **315** | 6,949,599,597 | 12,187,168 | 1,305,054 | 309,827 | 1,028 | 684,170,427 | 659,572 | 520,142 | 148,443 |
| **319** | 6,984,066,868 | 12,396,808 | 1,352,845 | 342,885 | 1,114 | 742,215,671 | 144,779 | 593,845 | 168,627 |
| **320** | 7,501,417,624 | 12,965,323 | 1,325,747 | 337,799 | 1,102 | 725,591,569 | 889,385 | 556,011 | 166,270 |
| **321** | 8,541,115,749 | 14,587,332 | 1,751,299 | 451,148 | 1,027 | 942,741,816 | 659,617 | 699,588 | 189,670 |
| **324** | 5,304,775,998 | 13,853,045 | 717,695 | 140,680 | 890 | 341,757,754 | 538,303 | 325,162 | 56,380 |
| **326** | 21,965,269,770 | 33,849,158 | 3,428,669 | 841,029 | 1,150 | 1,867,599,514 | 266,735 | 1,291,360 | 448,166 |
| **329** | 16,064,549,937 | 22,926,879 | 2,629,046 | 598,340 | 1,064 | 1,381,208,536 | 157,398 | 859,230 | 263,064 |
| **330** | 21,005,691,360 | 28,029,489 | 3,151,195 | 724,155 | 1,115 | 1,675,633,057 | 264,112 | 996,361 | 360,822 |

**Table S3**

List of genes involved in N-acquisition and transporters of AA-related compounds.

| Gene | Function | KO |
|---|---|---|
| ***Urease*** | | |
| ureA | urease subunit gamma | K01430 |
| ureB | urease subunit beta | K01429 |
| ureC | urease subunit alpha | K01428 |
| ureD, ureH | UreD-UreF-UreD complex; Hydolysies urea into ammonia and carbamic acid | K03190 |
| ureE | Hydolyses of urea into ammonia and carbamic acid | K03187 |
| ureF | UreD-UreF-UreD complex; Hydolysies urea into ammonia and carbamic acid | K03188 |
| ureG | UreD-UreF-UreD complex; Hydolyses of urea into ammonia and carbamic acid | K03189 |
| ***Urea transporter*** | | |
| urtA | urea transport system substrate-binding protein | K11959 |
| urtB | urea transport system permease protein | K11960 |
| urtC | urea transport system permease protein | K11961 |
| urtD | urea transport system ATP-binding protein | K11962 |
| urtE | urea transport system ATP-binding protein | K11963 |
| ***Assimilatory nitrate reduction, nitrate => nitrite*** | | |
| narB | ferredoxin-nitrate reductase [EC:1.7.7.2] | K00367 |
| NR | nitrate reductase (NAD(P)H) | K10534 |
| nasA | assimilatory nitrate reductase catalytic subunit | K00372 |
| nasB | assimilatory nitrate reductase electron transfer subunit | K00360 |
| narB | ferredoxin-nitrate reductase | K00367 |
| ***Assimilatory nitrite reduction, nitrite => ammonia*** | | |
| nirA | ferredoxin-nitrite reductase | K00366 |
| NIT-6 | nitrite reductase (NAD(P)H) | K17877 |
| ***Dissimilatory nitrate reductase, nitrate => nitrite*** | | |
| narG, narZ, nxrA | nitrate reductase / nitrite oxidoreductase, alpha subunit | K00370 |
| narH, narY, naxB | nitrate reductase / nitrite oxidoreductase, beta subunit | K00371 |
| narI, narV | nitrate reductase gamma subunit | K00374 |
| napA | periplasmic nitrate reductase NapA | K02567 |
| napB | cytochrome c-type protein NapB | K02568 |
| ***Disssimilatory nitrate reduction, nitrate => ammonia*** | | |
| nirD | nitrite reductase (NADH) small subunit | K00363 |
| nrfH | cytochrome c nitrite reductase small subunit | K15876 |
| nrfA | nitrite reductase (cytochrome c-552) | K03385 |
| ***Nitrogen fixation*** | | |
| nifA | nif-specific regulatory protein | K02584 |
| nifB | nitrogen fixation protein nifB | K02585 |
| nifD | nitrogenase molybdenum-iron protein alpha chain | K02586 |

| | | |
|---|---|---|
| nifE | nitrogenase molybdenum-iron cofactor synthesis protein | K02587 |
| nifH | nitrogenase iron protein | K02588 |
| nifK | nitrogenase molybdenum-iron protein beta chain | K02591 |
| nifN | nitrogen fixation protein nifN | K02592 |
| nifQ | nitrogen fixation protein nifQ | K15790 |
| nifT | nitrogen fixation protein nifT | K02593 |
| nifV | homocitrate synthase | K02594 |
| nifW | nitrogenase-stabilizing/protective protein | K02595 |
| nifX | nitrogen fixation protein NifX | K02596 |
| nifZ | nitrogen fixation protein NifZ | K02597 |
| vnfH | vanadium nitrogenase iron protein | K22899 |
| anfG | nitrogenase delta subunit | K00531 |
| vnfD | vanadium-dependent nitrogenase alpha chain | K22896 |
| vnfK | vanadium-dependent nitrogenase beta chain | K22897 |
| vnfG | vanadium nitrogenase delta subunit | K22898 |
| ***Cyanate lyase*** | | |
| cynR | cyn operon transcriptional activator | K11921 |
| cynS | cyanate lyase | K01725 |
| cynT, can | carbonic anhydrase | K01673 |
| ***Ammonium transporter*** | | |
| amt, AMT, MEP | ammonium transporter, Amt family | K03320 |
| ***Nitrilase*** | | |
| | nitrilase [EC:3.5.1.49] | K01501 |
| ***Formamidase*** | | |
| | formamidase [EC:3.5.1.49] | K01455 |
| ***Gluamate Synthase*** | | |
| gdhA | glutamate dehydrogenase (NAD(P)+) [EC:1.4.1.3] | K00261 |
| gdhA | glutamate dehydrogenase (NADP+) [EC:1.4.1.4] | K00262 |
| GDH2 | glutamate dehydrogenase [EC:1.4.1.2] | K15371 |
| glnA | glutamine synthetase [EC:6.3.1.2] | K01915 |
| gltB | glutamate synthase (NADPH) large chain [EC:1.4.1.13] | K00265 |
| gltD | glutamate synthase (NADPH) small chain [EC:1.4.1.13] | K00266 |
| gltS | glutamate synthase (ferredoxin) [EC:1.4.7.1] | K00284 |
| ***Denitrification*** | | |
| nirK | nitrite reductase (NO-forming) [EC:1.7.2.1] | K00368 |
| nirS | nitrite reductase (NO-forming) / hydroxylamine reductase [EC:1.7.2.1 1.7.99.1] | K15864 |
| norB | nitric oxide reductase subunit B [EC:1.7.2.5] | K04561 |
| norC | nitric oxide reductase subunit C | K02305 |
| nosZ | nitrous-oxide reductase [EC:1.7.2.4] | K00376 |
| ***Arginine transporter*** | | |

| | | |
|---|---|---|
| artJ | arginine transport system | K09996 |
| artI | arginine transport system substrate-binding protein | K09997 |
| artM | arginine transport system permease protein | K09998 |
| artQ | arginine transport system permease protein | K09999 |
| artP | arginine transport system ATP-binding protein [EC:7.4.2.1] | K10000 |

***Arginine/Ornithine transporter***

| | | |
|---|---|---|
| aotJ | arginine/ornithine transport system substrate-binding protein | K10022 |
| aotM | arginine/ornithine transport system permease protein | K10023 |
| aotQ | arginine/ornithine transport system permease protein | K10024 |
| aotP | arginine/ornithine transport system ATP-binding protein [EC:7.4.2.1] | K10025 |

***Arginine/Lysine/Histidine transporter***

| | | |
|---|---|---|
| artR | arginine/lysine/histidine transport system ATP-binding protein [EC:7.4.2.1] | K23060 |
| artP | arginine/lysine/histidine transporter system substrate-binding protein | K23059 |
| artQ | arginine/lysine/histidine transport system permease protein | K17077 |

***Arginine/Lysine/Histidine/Glutamate transporter***

| | | |
|---|---|---|
| bgtB | arginine/lysine/histidine/glutamine transport system substrate-binding and permease protein | K17062 |
| bgtA | arginine/lysine/histidine/glutamine transport system ATP-binding protein [EC:7.4.2.1] | K17063 |

***Aspartate/Glutamate/Glutamine transporter***

| | | |
|---|---|---|
| peb1A, glnH | aspartate/glutamate/glutamine transport system substrate-binding protein | K10039 |
| peb1B, glnP, glnM | aspartate/glutamate/glutamine transport system permease protein | K10040 |
| peb1C, glnQ | aspartate/glutamate/glutamine transport system ATP-binding protein [EC:7.4.2.1] | K10041 |

***Branched chain amino acids transporter***

| | | |
|---|---|---|
| livK | branched-chain amino acid transport system substrate-binding protein | K01999 |
| livH | branched-chain amino acid transport system permease protein | K01997 |
| livM | branched-chain amino acid transport system permease protein | K01998 |
| livG | branched-chain amino acid transport system ATP-binding protein | K01995 |
| livF | branched-chain amino acid transport system ATP-binding protein | K01996 |

***Cysteine transporter***

| | | |
|---|---|---|
| fliY, tcyA | L-cystine transport system substrate-binding protein | K02424 |
| tcyB, yecS | L-cystine transport system permease protein | K10009 |
| tcyC, yecC | L-cystine transport system ATP-binding protein [EC:7.4.2.1] | K10010 |
| tcyJ | L-cystine transport system substrate-binding protein | K16956 |
| tcyK | L-cystine transport system substrate-binding protein | K16957 |
| tcyL | L-cystine transport system permease protein | K16958 |
| tcyM | L-cystine transport system permease protein | K16959 |
| tcyN | L-cystine transport system ATP-binding protein [EC:7.4.2.1] | K16960 |

***Dipeptide transporter***

| | | |
|---|---|---|
| dppE | dipeptide transport system substrate-binding protein | K16199 |
| dppB | dipeptide transport system permease protein | K16200 |
| dppC | dipeptide transport system permease protein | K16201 |
| dppD | dipeptide transport system ATP-binding protein | K16202 |
| dppA | dipeptide transport system substrate-binding protein | K12368 |
| dppF | dipeptide transport system ATP-binding protein | K12372 |
| ***General amino acid transporter*** | | |
| aapJ, bztA | general L-amino acid transport system substrate-binding protein | K09969 |
| aapQ, bztB | general L-amino acid transport system permease protein | K09970 |
| aapM, bztC | general L-amino acid transport system permease protein | K09971 |
| aapP, bztD | general L-amino acid transport system ATP-binding protein [EC:7.4.2.1] | K09972 |
| ***Glutamate transporter*** | | |
| gluB | glutamate transport system substrate-binding protein | K10005 |
| gluC | glutamate transport system permease protein | K10006 |
| gluD | glutamate transport system permease protein | K10007 |
| gluA | glutamate transport system ATP-binding protein [EC:7.4.2.1] | K10008 |
| ***Glutamate/Aspartate transporter*** | | |
| gltI, aatJ | glutamate/aspartate transport system substrate-binding protein | K10001 |
| gltK, aatM | glutamate/aspartate transport system permease protein | K10002 |
| gltJ, aatQ | glutamate/aspartate transport system permease protein | K10003 |
| gltL, aatP | glutamate/aspartate transport system ATP-binding protein [EC:7.4.2.1] | K10004 |
| ***Glutamine transporter*** | | |
| glnH | glutamine transport system substrate-binding protein | K10036 |
| glnP | glutamine transport system permease protein | K10037 |
| glnQ | glutamine transport system ATP-binding protein [EC:7.4.2.1] | K10038 |
| ***Gluthadione transporter*** | | |
| gsiB | glutathione transport system substrate-binding protein | K13889 |
| gsiC | glutathione transport system permease protein | K13890 |
| gsiD | glutathione transport system permease protein | K13891 |
| gsiA | glutathione transport system ATP-binding protein | K13892 |
| ***Glycine/Proline transporter*** | | |
| proX | glycine betaine/proline transport system substrate-binding protein | K02002 |
| proW | glycine betaine/proline transport system permease protein | K02001 |
| proV | glycine betaine/proline transport system ATP-binding protein [EC:3.6.3.32] | K02000 |
| ***Histidine transporter*** | | |
| hisJ | histidine transport system substrate-binding protein | K10014 |
| hisM | histidine transport system permease protein | K10015 |
| hisQ | histidine transport system permease protein | K10016 |

| | | |
|---|---|---|
| hisP | histidine transport system ATP-binding protein [EC:7.4.2.1] | K10017 |
| ***Hydroxyproline transporter*** | | |
| lhpP | hydroxyproline transporter system substrate-binding protein | K23061 |
| lhpM | hydroxyproline transport system permease protein | K23062 |
| lhpN | hydroxyproline transport system permease protein | K23063 |
| lhpO | hydroxyproline transport system ATP-binding protein [EC:7.4.2.1] | K23064 |
| ***Lysine transporter*** | | |
| lysX1 | putative lysine transport system substrate-binding protein | K17073 |
| lysX2 | putative lysine transport system permease protein | K17074 |
| lysY | putative lysine transport system ATP-binding protein [EC:3.6.3.-] | K17076 |
| ***Lysine/Arginine transporter*** | | |
| argT | lysine/arginine/ornithine transport system substrate-binding protein | K10013 |
| hisM | histidine transport system permease protein | K10015 |
| hisQ | histidine transport system permease protein | K10016 |
| hisP | histidine transport system ATP-binding protein [EC:7.4.2.1] | K10017 |
| ***Lysine/Arginine/Ornithine/Histidine/Octopine transporter*** | | |
| PA5153 | Putative lysine/arginine/ornithine/histidine/octopine transport system | K23055 |
| PA5154 | putative lysine/arginine/ornithine/histidine/octopine transport system permease protein | K23056 |
| PA5155 | putative lysine/arginine/ornithine/histidine/octopine transport system permease protein | K23057 |
| PA5152 | putative lysine/arginine/ornithine/histidine/octopine transport system ATP-binding protein [EC:7.4.2.1] | K23058 |
| ***Methionine transporter*** | | |
| metQ | D-methionine transport system substrate-binding protein | K02073 |
| metI | D-methionine transport system permease protein | K02072 |
| netN | D-methionine transport system ATP-binding protein | K02071 |
| ***S-Methylcysteine transporter*** | | |
| yxeM | putative S-methylcysteine transport system substrate-binding protein | K16961 |
| yxeN | putative S-methylcysteine transport system permease protein | K16962 |
| yxeO | putative S-methylcysteine transport system ATP-binding protein | K16963 |
| ***neutral amino acid/Histidine transporter*** | | |
| natB | neutral amino acid transport system substrate-binding protein | K11954 |
| natC | neutral amino acid transport system permease protein | K11955 |
| natD | neutral amino acid transport system permease protein | K11956 |
| natA | neutral amino acid transport system ATP-binding protein | K11957 |
| natE | neutral amino acid transport system ATP-binding protein | K11958 |
| ***Oligopeptide transporter*** | | |
| oppA, mppA | oligopeptide transport system substrate-binding protein | K15580 |

| | | |
|---|---|---|
| oppB | oligopeptide transport system permease protein | K15581 |
| oppC | oligopeptide transport system permease protein | K15582 |
| oppD | oligopeptide transport system ATP-binding protein | K15583 |
| oppF | oligopeptide transport system ATP-binding protein | K10823 |

***Octopine/Nopaline transporter***

| | | |
|---|---|---|
| occT, nocT | octopine/nopaline transport system substrate-binding protein | K10018 |
| occM, nocM | octopine/nopaline transport system permease protein | K10019 |
| occQ, nocQ | octopine/nopaline transport system permease protein | K10020 |
| occP, nocP | octopine/nopaline transport system ATP-binding protein [EC:7.4.2.1] | K10021 |

### Supplementary Figures

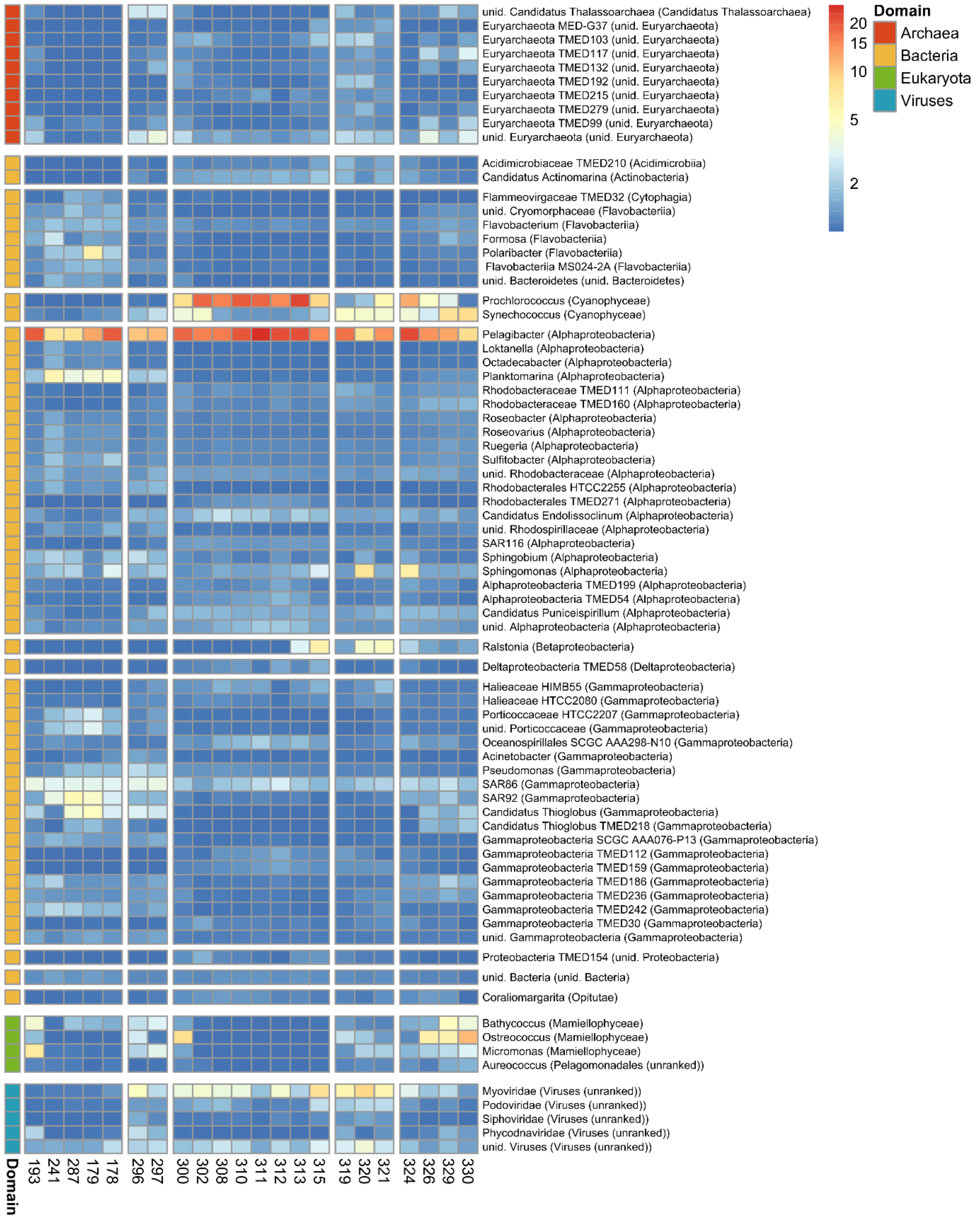

### Figure S1

Relative abundance of prominent prokaryotic and eukaryotic genera and viral families along the Atlantic Ocean transect at stations 193 to 330 (for station details see Table S1).

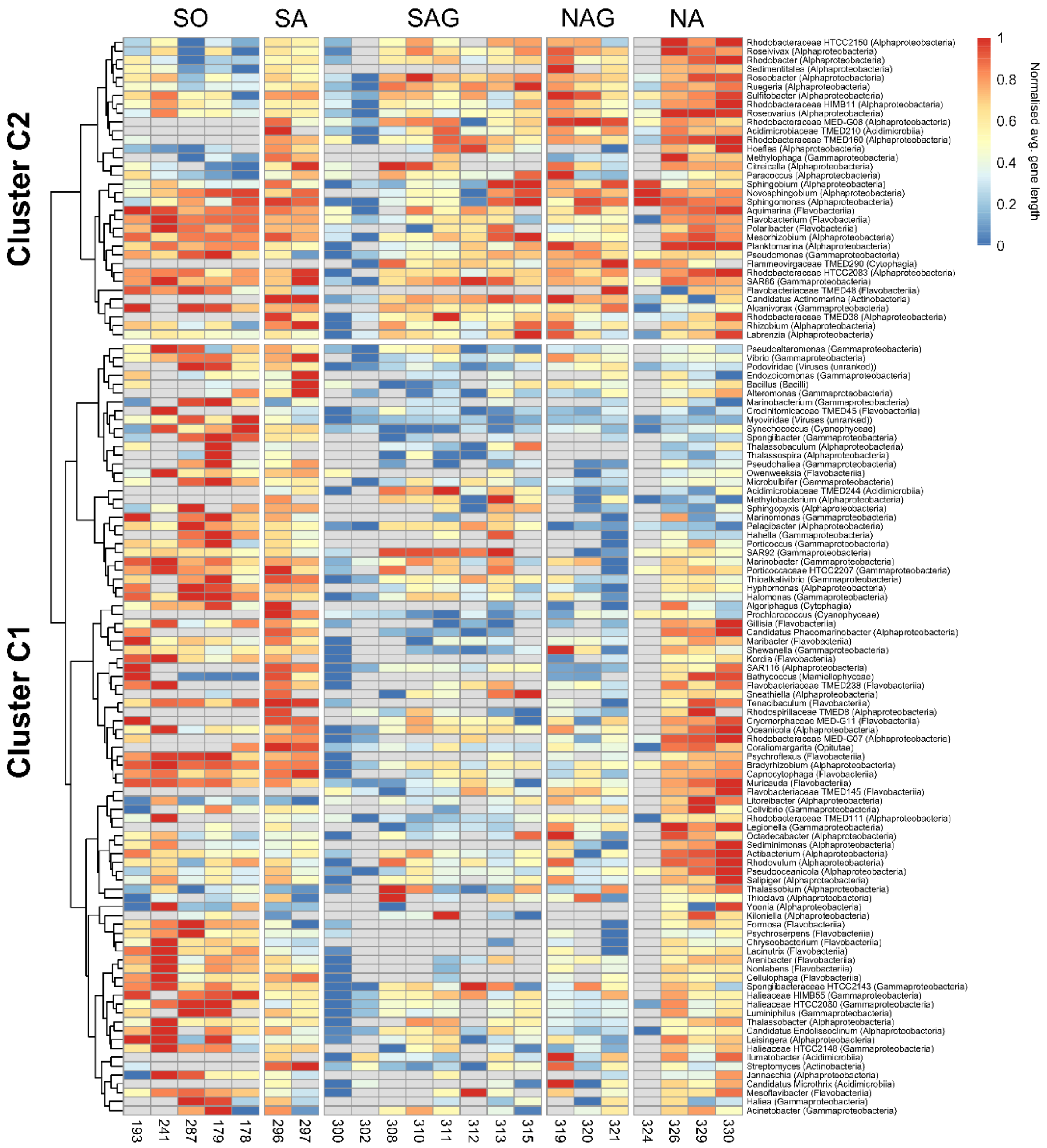

**Figure S2**

Normalized AGL distribution among prokaryotic and eukaryotic genera as well as viral families along the Atlantic Ocean transect between 62° and 47°N at stations 193 to 330 (for station details see Table S1). Only taxa with ≥ 50 complete genes (~10kb) at ≥50% of the stations were considered. Euclidean distances were calculated and clustered using Ward.D2 clustering.

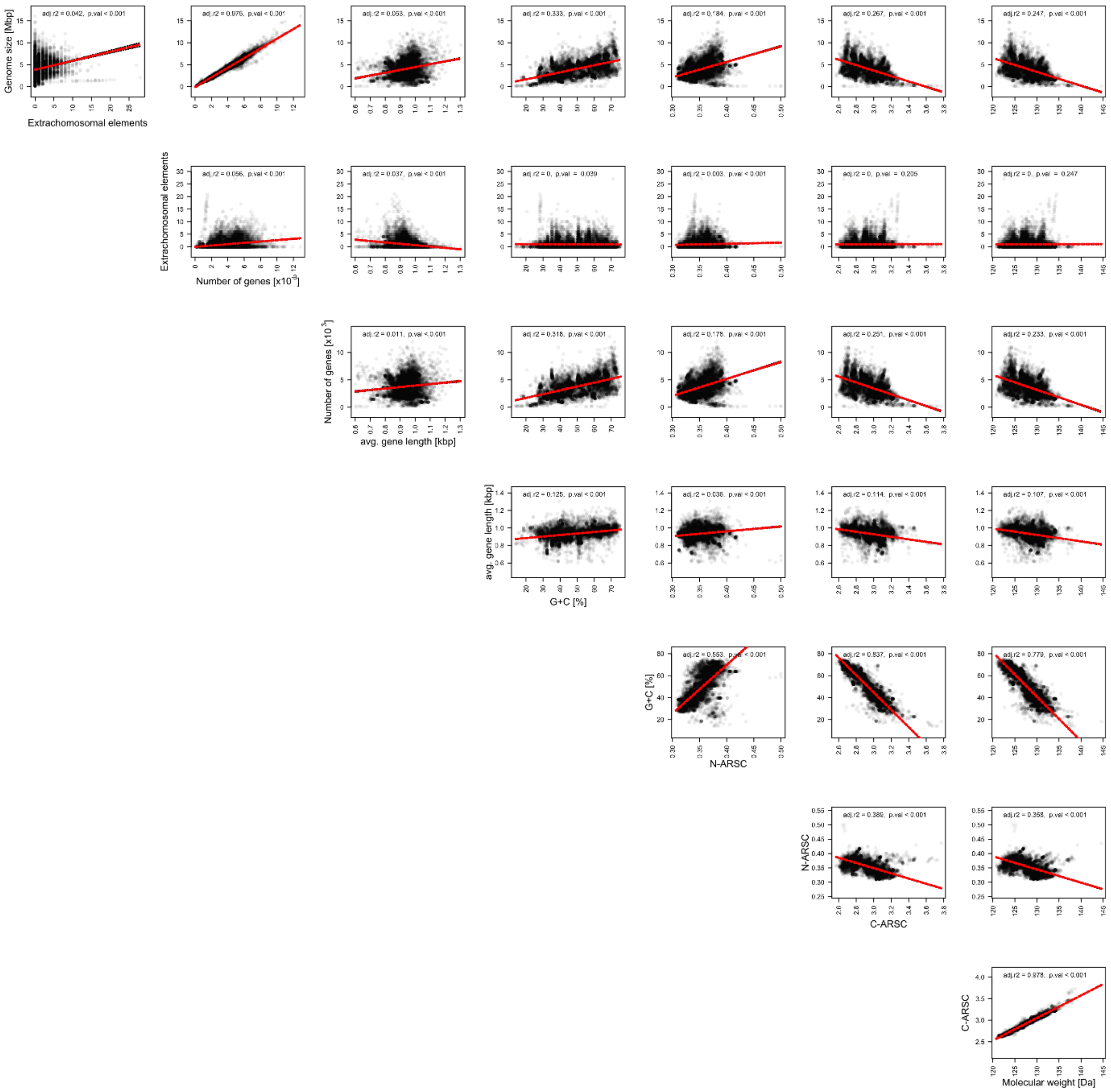

**Figure S3**

Correlation matrix of genome size, extrachromosomal elements), number of genes, AGL, G+C-content, N-ARSC and C-ARSC of all prokaryotic genomes available at GeneBank in January 2020

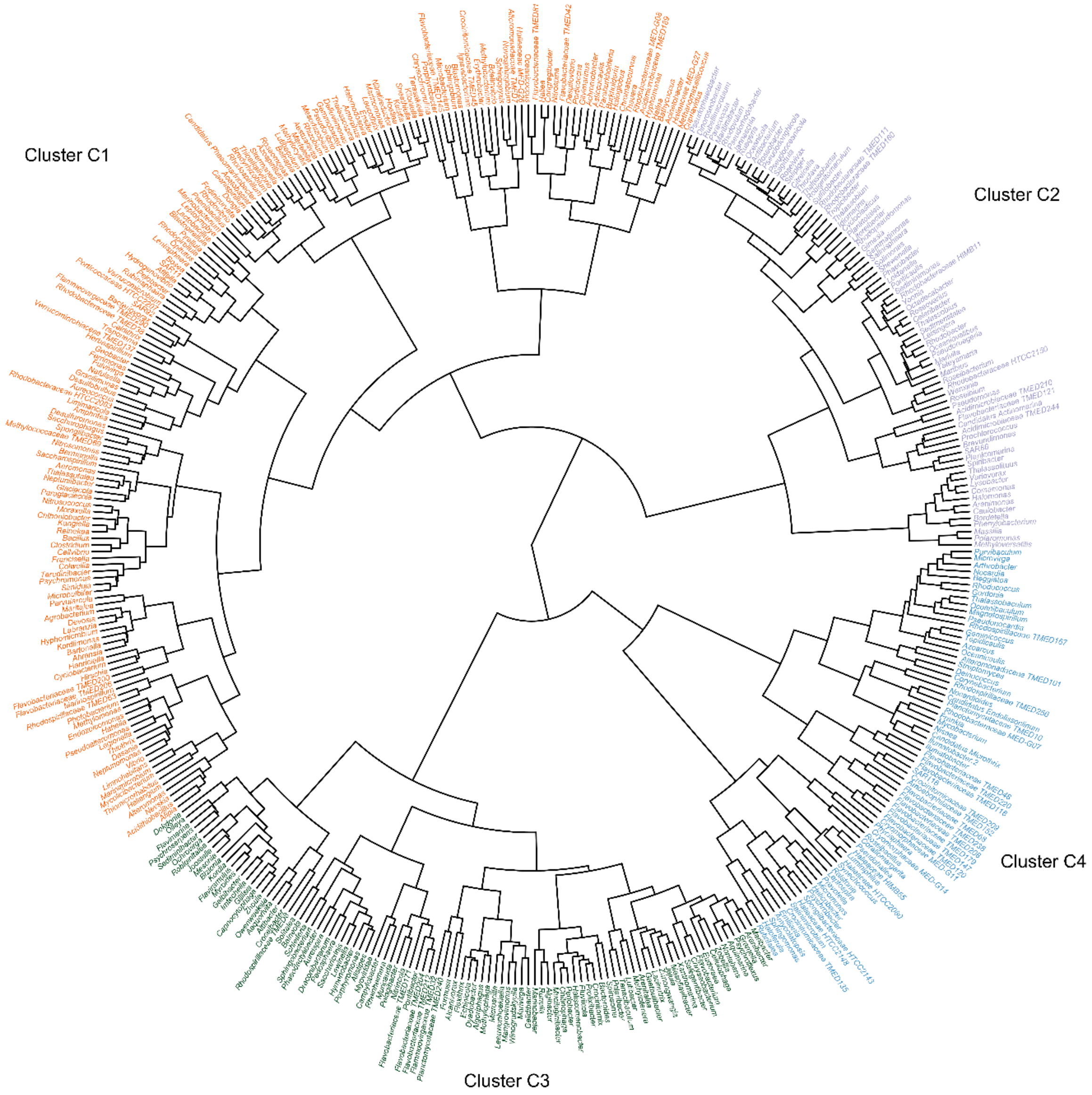

**Figure S4**

Clustered distribution of the genomic G+C content of the AOM. Only taxa with ≥ 50 complete genes (~10kb) at >50% of stations were considered. Euclidean distances were calculated and clustered using Ward.D2 clustering.

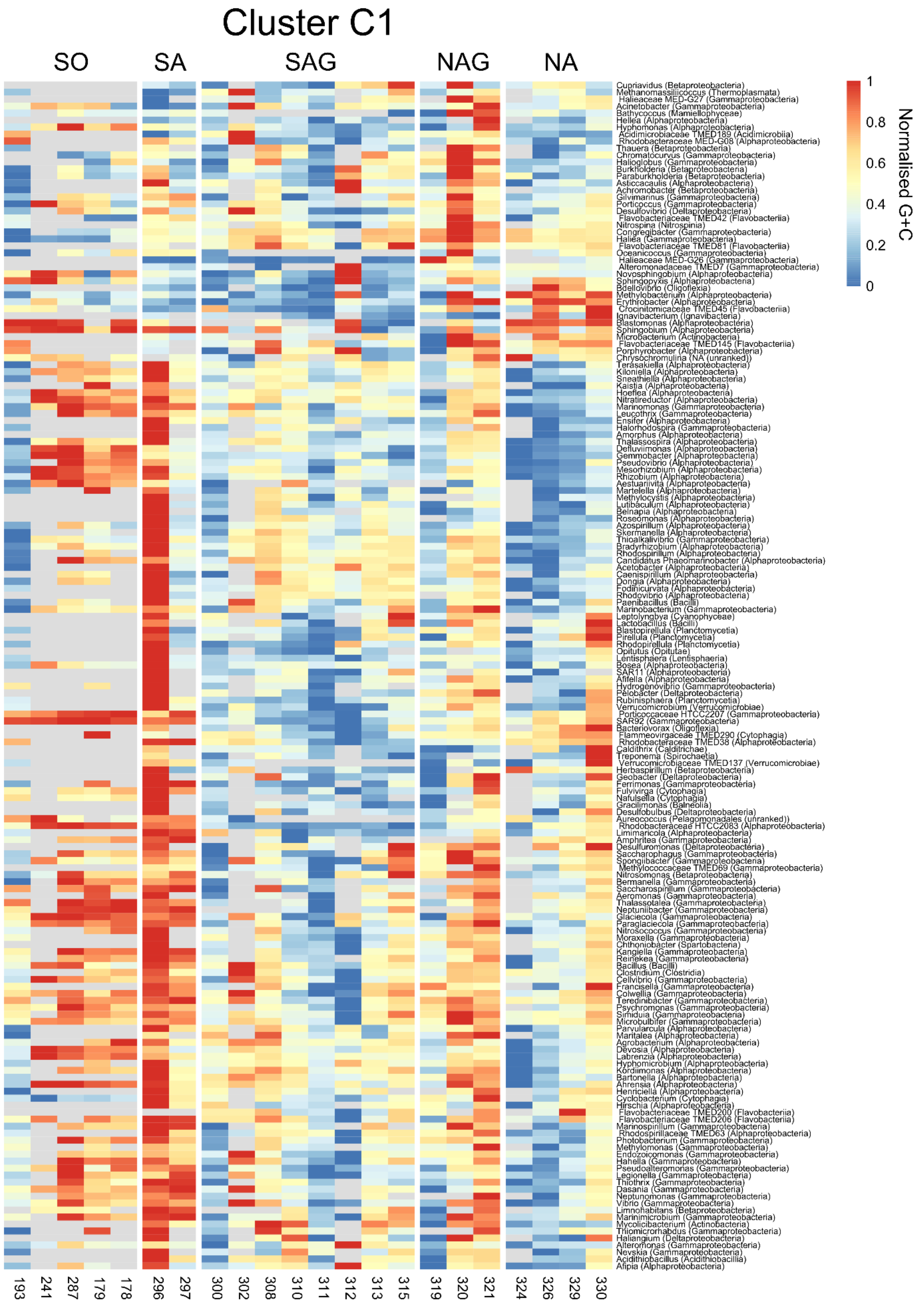

Cluster C1
SO
SA
SAG
NAG
NA
Cupriavidus (Betaproteobacteria)
Methanomassiliicoccus (Thermoplasmata)
Halieaceae MED-G27 (Gammaproteobacteria)
Acinetobacter (Gammaproteobacteria)
Bathycoccus (Mamiellophyceae)
Hellea (Alphaproteobacteria)
Hyphomonas (Alphaproteobacteria)
Acidimicrobiaceae TMED189 (Acidimicrobiia)
Rhodobacteraceae MED-G08 (Alphaproteobacteria)
Thauera (Betaproteobacteria)
Chromatocurvus (Gammaproteobacteria)
Halioglobus (Gammaproteobacteria)
Burkholderia (Betaproteobacteria)
Paraburkholderia (Betaproteobacteria)
Asticcacaulis (Alphaproteobacteria)
Achromobacter (Betaproteobacteria)
Gilvimarinus (Gammaproteobacteria)
Porticoccus (Gammaproteobacteria)
Desulfovibrio (Deltaproteobacteria)
Flavobacteriaceae TMED42 (Flavobacteriia)
Nitrospina (Nitrospinia)
Congregibacter (Gammaproteobacteria)
Haliea (Gammaproteobacteria)
Flavobacteriaceae TMED81 (Flavobacteriia)
Oceanicoccus (Gammaproteobacteria)
Halieaceae MED-G26 (Gammaproteobacteria)
Alteromonadaceae TMED7 (Gammaproteobacteria)
Novosphingobium (Alphaproteobacteria)
Sphingopyxis (Alphaproteobacteria)
Bdellovibrio (Oligoflexia)
Methylobacterium (Alphaproteobacteria)
Erythrobacter (Alphaproteobacteria)
Crocinitomicaceae TMED45 (Flavobacteriia)
Ignavibacterium (Ignavibacteria)
Blastomonas (Alphaproteobacteria)
Sphingobium (Alphaproteobacteria)
Microbacterium (Actinobacteria)
Flavobacteriaceae TMED145 (Flavobacteriia)
Porphyrobacter (Alphaproteobacteria)
Chrysochromulina (NA (unranked))
Terasakiella (Alphaproteobacteria)
Kiloniella (Alphaproteobacteria)
Sneathiella (Alphaproteobacteria)
Kaistia (Alphaproteobacteria)
Hoeflea (Alphaproteobacteria)
Nitratireductor (Alphaproteobacteria)
Marinomonas (Gammaproteobacteria)
Leucothrix (Gammaproteobacteria)
Ensifer (Alphaproteobacteria)
Halorhodospira (Gammaproteobacteria)
Amorphus (Alphaproteobacteria)
Thalassospira (Alphaproteobacteria)
Defluviimonas (Alphaproteobacteria)
Gemmobacter (Alphaproteobacteria)
Pseudovibrio (Alphaproteobacteria)
Mesorhizobium (Alphaproteobacteria)
Rhizobium (Alphaproteobacteria)
Aestuariivita (Alphaproteobacteria)
Martelella (Alphaproteobacteria)
Methylocystis (Alphaproteobacteria)
Lutibaculum (Alphaproteobacteria)
Belnapia (Alphaproteobacteria)
Roseomonas (Alphaproteobacteria)
Azospirillum (Alphaproteobacteria)
Skermanella (Alphaproteobacteria)
Thioalkalivibrio (Gammaproteobacteria)
Bradyrhizobium (Alphaproteobacteria)
Rhodospirillum (Alphaproteobacteria)
Candidatus Phaeomarinobacter (Alphaproteobacteria)
Acetobacter (Alphaproteobacteria)
Caenispirillum (Alphaproteobacteria)
Dongia (Alphaproteobacteria)
Fodinicurvata (Alphaproteobacteria)
Rhodovibrio (Alphaproteobacteria)
Paenibacillus (Bacilli)
Marinobacterium (Gammaproteobacteria)
Leptolyngbya (Cyanophyceae)
Lactobacillus (Bacilli)
Blastopirellula (Planctomycetia)
Pirellula (Planctomycetia)
Rhodopirellula (Planctomycetia)
Opitutus (Opitutae)
Lentisphaera (Lentisphaeria)
Bosea (Alphaproteobacteria)
SAR11 (Alphaproteobacteria)
Afifella (Alphaproteobacteria)
Hydrogenovibrio (Gammaproteobacteria)
Pelobacter (Deltaproteobacteria)
Rubinisphaera (Planctomycetia)
Verrucomicrobium (Verrucomicrobiae)
Porticoccaceae HTCC2207 (Gammaproteobacteria)
SAR92 (Gammaproteobacteria)
Bacteriovorax (Oligoflexia)
Flammeovirgaceae TMED290 (Cytophagia)
Rhodobacteraceae TMED38 (Alphaproteobacteria)
Caldithrix (Calditrichae)
Treponema (Spirochaetia)
Verrucomicrobiaceae TMED137 (Verrucomicrobiae)
Herbaspirillum (Betaproteobacteria)
Geobacter (Deltaproteobacteria)
Ferrimonas (Gammaproteobacteria)
Fulvivirga (Cytophagia)
Nafulsella (Cytophagia)
Gracilimonas (Balneolia)
Desulfobulbus (Deltaproteobacteria)
Aureococcus (Pelagomonadales (unranked))
Rhodobacteraceae HTCC2083 (Alphaproteobacteria)
Limimaricola (Alphaproteobacteria)
Amphritea (Gammaproteobacteria)
Desulfuromonas (Deltaproteobacteria)
Saccharophagus (Gammaproteobacteria)
Spongiibacter (Gammaproteobacteria)
Methylococcaceae TMED69 (Gammaproteobacteria)
Nitrosomonas (Betaproteobacteria)
Bermanella (Gammaproteobacteria)
Saccharospirillum (Gammaproteobacteria)
Aeromonas (Gammaproteobacteria)
Thalassotalea (Gammaproteobacteria)
Neptuniibacter (Gammaproteobacteria)
Glaciecola (Gammaproteobacteria)
Paraglaciecola (Gammaproteobacteria)
Nitrosococcus (Gammaproteobacteria)
Moraxella (Gammaproteobacteria)
Chthoniobacter (Spartobacteria)
Kangiella (Gammaproteobacteria)
Reinekea (Gammaproteobacteria)
Bacillus (Bacilli)
Clostridium (Clostridia)
Cellvibrio (Gammaproteobacteria)
Francisella (Gammaproteobacteria)
Colwellia (Gammaproteobacteria)
Teredinibacter (Gammaproteobacteria)
Psychromonas (Gammaproteobacteria)
Simiduia (Gammaproteobacteria)
Microbulbifer (Gammaproteobacteria)
Parvularcula (Alphaproteobacteria)
Maritalea (Alphaproteobacteria)
Agrobacterium (Alphaproteobacteria)
Devosia (Alphaproteobacteria)
Labrenzia (Alphaproteobacteria)
Hyphomicrobium (Alphaproteobacteria)
Kordiimonas (Alphaproteobacteria)
Bartonella (Alphaproteobacteria)
Ahrensia (Alphaproteobacteria)
Henriciella (Alphaproteobacteria)
Cyclobacterium (Cytophagia)
Hirschia (Alphaproteobacteria)
Flavobacteriaceae TMED200 (Flavobacteriia)
Flavobacteriaceae TMED206 (Flavobacteriia)
Marinospirillum (Gammaproteobacteria)
Rhodospirillaceae TMED63 (Alphaproteobacteria)
Photobacterium (Gammaproteobacteria)
Methylomonas (Gammaproteobacteria)
Endozoicomonas (Gammaproteobacteria)
Hahella (Gammaproteobacteria)
Pseudoalteromonas (Gammaproteobacteria)
Legionella (Gammaproteobacteria)
Thiothrix (Gammaproteobacteria)
Dasania (Gammaproteobacteria)
Neptunomonas (Gammaproteobacteria)
Vibrio (Gammaproteobacteria)
Limnohabitans (Betaproteobacteria)
Marinimicrobium (Gammaproteobacteria)
Mycolicibacterium (Actinobacteria)
Thiomicrorhabdus (Gammaproteobacteria)
Haliangium (Deltaproteobacteria)
Alteromonas (Gammaproteobacteria)
Nevskia (Gammaproteobacteria)
Acidithiobacillus (Acidithiobacillia)
Afipia (Alphaproteobacteria)
193
241
287
179
178
296
297
300
302
308
310
311
312
313
315
319
320
321
324
326
329
330
1
0.8
0.6
0.4
0.2
0
Normalised G+C

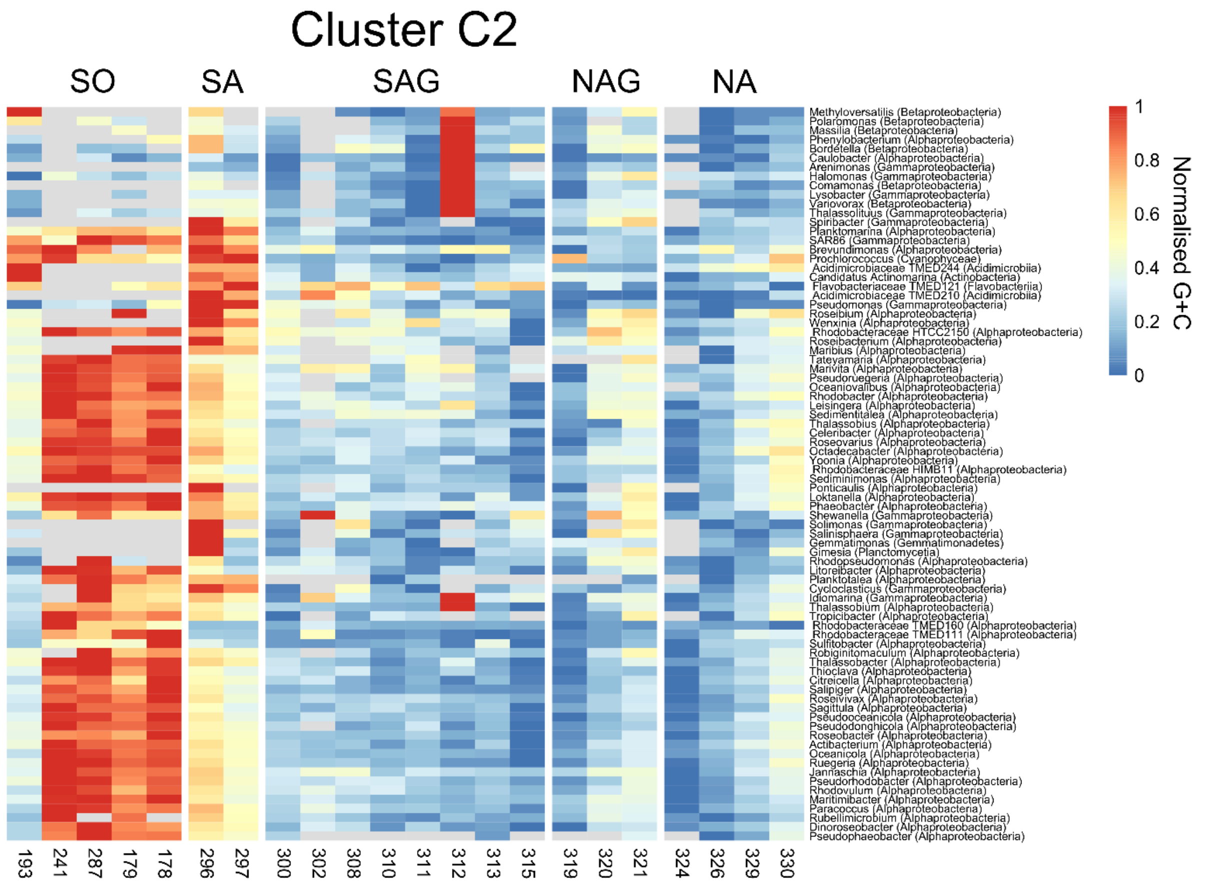

Cluster C2
SO
SA
SAG
NAG
NA
Methyloversatilis (Betaproteobacteria)
Polaromonas (Betaproteobacteria)
Massilia (Betaproteobacteria)
Phenylobacterium (Alphaproteobacteria)
Bordetella (Betaproteobacteria)
Caulobacter (Alphaproteobacteria)
Arenimonas (Gammaproteobacteria)
Halomonas (Gammaproteobacteria)
Comamonas (Betaproteobacteria)
Lysobacter (Gammaproteobacteria)
Variovorax (Betaproteobacteria)
Thalassolituus (Gammaproteobacteria)
Spiribacter (Gammaproteobacteria)
Planktomarina (Alphaproteobacteria)
SAR86 (Gammaproteobacteria)
Brevundimonas (Alphaproteobacteria)
Prochlorococcus (Cyanophyceae)
Acidimicrobiaceae TMED244 (Acidimicrobiia)
Candidatus Actinomarina (Actinobacteria)
Flavobacteriaceae TMED121 (Flavobacteriia)
Acidimicrobiaceae TMED210 (Acidimicrobiia)
Pseudomonas (Gammaproteobacteria)
Roseibium (Alphaproteobacteria)
Wenxinia (Alphaproteobacteria)
Rhodobacteraceae HTCC2150 (Alphaproteobacteria)
Roseibacterium (Alphaproteobacteria)
Maribius (Alphaproteobacteria)
Tateyamaria (Alphaproteobacteria)
Marivita (Alphaproteobacteria)
Pseudoruegeria (Alphaproteobacteria)
Oceaniovalibus (Alphaproteobacteria)
Rhodobacter (Alphaproteobacteria)
Leisingera (Alphaproteobacteria)
Sedimentitalea (Alphaproteobacteria)
Thalassobius (Alphaproteobacteria)
Celeribacter (Alphaproteobacteria)
Roseovarius (Alphaproteobacteria)
Octadecabacter (Alphaproteobacteria)
Yoonia (Alphaproteobacteria)
Rhodobacteraceae HIMB11 (Alphaproteobacteria)
Sediminimonas (Alphaproteobacteria)
Ponticaulis (Alphaproteobacteria)
Loktanella (Alphaproteobacteria)
Phaeobacter (Alphaproteobacteria)
Shewanella (Gammaproteobacteria)
Solimonas (Gammaproteobacteria)
Salinisphaera (Gammaproteobacteria)
Gemmatimonas (Gemmatimonadetes)
Gimesia (Planctomycetia)
Rhodopseudomonas (Alphaproteobacteria)
Litoreibacter (Alphaproteobacteria)
Planktotalea (Alphaproteobacteria)
Cycloclasticus (Gammaproteobacteria)
Idiomarina (Gammaproteobacteria)
Thalassobium (Alphaproteobacteria)
Tropicibacter (Alphaproteobacteria)
Rhodobacteraceae TMED160 (Alphaproteobacteria)
Rhodobacteraceae TMED111 (Alphaproteobacteria)
Sulfitobacter (Alphaproteobacteria)
Robiginitomaculum (Alphaproteobacteria)
Thalassobacter (Alphaproteobacteria)
Thioclava (Alphaproteobacteria)
Citreicella (Alphaproteobacteria)
Salipiger (Alphaproteobacteria)
Roseivivax (Alphaproteobacteria)
Sagittula (Alphaproteobacteria)
Pseudooceanicola (Alphaproteobacteria)
Pseudodonghicola (Alphaproteobacteria)
Roseobacter (Alphaproteobacteria)
Actibacterium (Alphaproteobacteria)
Oceanicola (Alphaproteobacteria)
Ruegeria (Alphaproteobacteria)
Jannaschia (Alphaproteobacteria)
Pseudorhodobacter (Alphaproteobacteria)
Rhodovulum (Alphaproteobacteria)
Maritimibacter (Alphaproteobacteria)
Paracoccus (Alphaproteobacteria)
Rubellimicrobium (Alphaproteobacteria)
Dinoroseobacter (Alphaproteobacteria)
Pseudophaeobacter (Alphaproteobacteria)
193
241
287
179
178
296
297
300
302
308
310
311
312
313
315
319
320
321
324
326
329
330
1
0.8
0.6
0.4
0.2
0
Normalised G+C

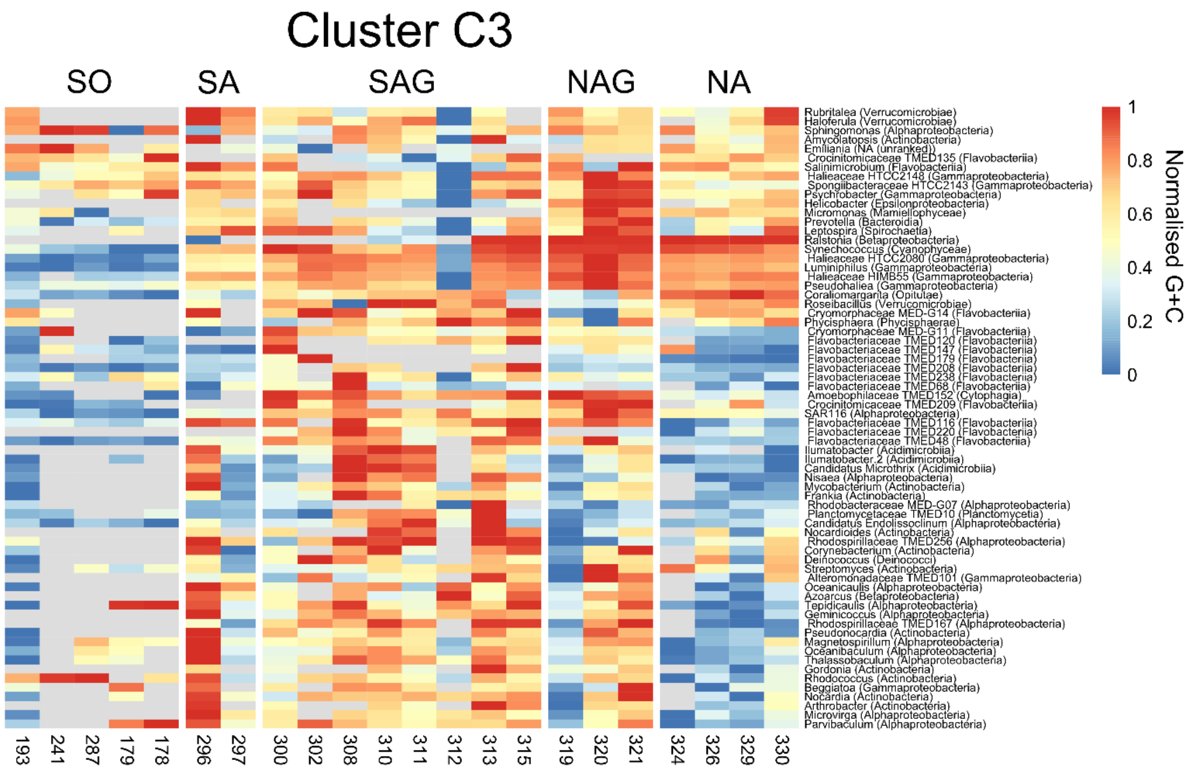

Cluster C3
SO
SA
SAG
NAG
NA
Rubritalea (Verrucomicrobiae)
Haloferula (Verrucomicrobiae)
Sphingomonas (Alphaproteobacteria)
Amycolatopsis (Actinobacteria)
Emiliania (NA (unranked))
Crocinitomicaceae TMED135 (Flavobacteriia)
Salinimicrobium (Flavobacteriia)
Halieaceae HTCC2148 (Gammaproteobacteria)
Spongiibacteraceae HTCC2143 (Gammaproteobacteria)
Psychrobacter (Gammaproteobacteria)
Helicobacter (Epsilonproteobacteria)
Micromonas (Mamiellophyceae)
Prevotella (Bacteroidia)
Leptospira (Spirochaetia)
Ralstonia (Betaproteobacteria)
Synechococcus (Cyanophyceae)
Halieaceae HTCC2080 (Gammaproteobacteria)
Luminiphilus (Gammaproteobacteria)
Halieaceae HIMB55 (Gammaproteobacteria)
Pseudohaliea (Gammaproteobacteria)
Coraliomargarita (Opitutae)
Roseibacillus (Verrucomicrobiae)
Cryomorphaceae MED-G14 (Flavobacteriia)
Phycisphaera (Phycisphaerae)
Cryomorphaceae MED-G11 (Flavobacteriia)
Flavobacteriaceae TMED120 (Flavobacteriia)
Flavobacteriaceae TMED147 (Flavobacteriia)
Flavobacteriaceae TMED179 (Flavobacteriia)
Flavobacteriaceae TMED208 (Flavobacteriia)
Flavobacteriaceae TMED238 (Flavobacteriia)
Flavobacteriaceae TMED68 (Flavobacteriia)
Amoebophilaceae TMED152 (Cytophagia)
Crocinitomicaceae TMED209 (Flavobacteriia)
SAR116 (Alphaproteobacteria)
Flavobacteriaceae TMED116 (Flavobacteriia)
Flavobacteriaceae TMED220 (Flavobacteriia)
Flavobacteriaceae TMED48 (Flavobacteriia)
Ilumatobacter (Acidimicrobiia)
Ilumatobacter.2 (Acidimicrobiia)
Candidatus Microthrix (Acidimicrobiia)
Nisaea (Alphaproteobacteria)
Mycobacterium (Actinobacteria)
Frankia (Actinobacteria)
Rhodobacteraceae MED-G07 (Alphaproteobacteria)
Planctomycetaceae TMED10 (Planctomycetia)
Candidatus Endolissoclinum (Alphaproteobacteria)
Nocardioides (Actinobacteria)
Rhodospirillaceae TMED256 (Alphaproteobacteria)
Corynebacterium (Actinobacteria)
Deinococcus (Deinococci)
Streptomyces (Actinobacteria)
Alteromonadaceae TMED101 (Gammaproteobacteria)
Oceanicaulis (Alphaproteobacteria)
Azoarcus (Betaproteobacteria)
Tepidicaulis (Alphaproteobacteria)
Geminicoccus (Alphaproteobacteria)
Rhodospirillaceae TMED167 (Alphaproteobacteria)
Pseudonocardia (Actinobacteria)
Magnetospirillum (Alphaproteobacteria)
Oceanibaculum (Alphaproteobacteria)
Thalassobaculum (Alphaproteobacteria)
Gordonia (Actinobacteria)
Rhodococcus (Actinobacteria)
Beggiatoa (Gammaproteobacteria)
Nocardia (Actinobacteria)
Arthrobacter (Actinobacteria)
Microvirga (Alphaproteobacteria)
Parvibaculum (Alphaproteobacteria)
193
241
287
179
178
296
297
300
302
308
310
311
312
313
315
319
320
321
324
326
329
330
1
0.8
0.6
0.4
0.2
0
Normalised G+C

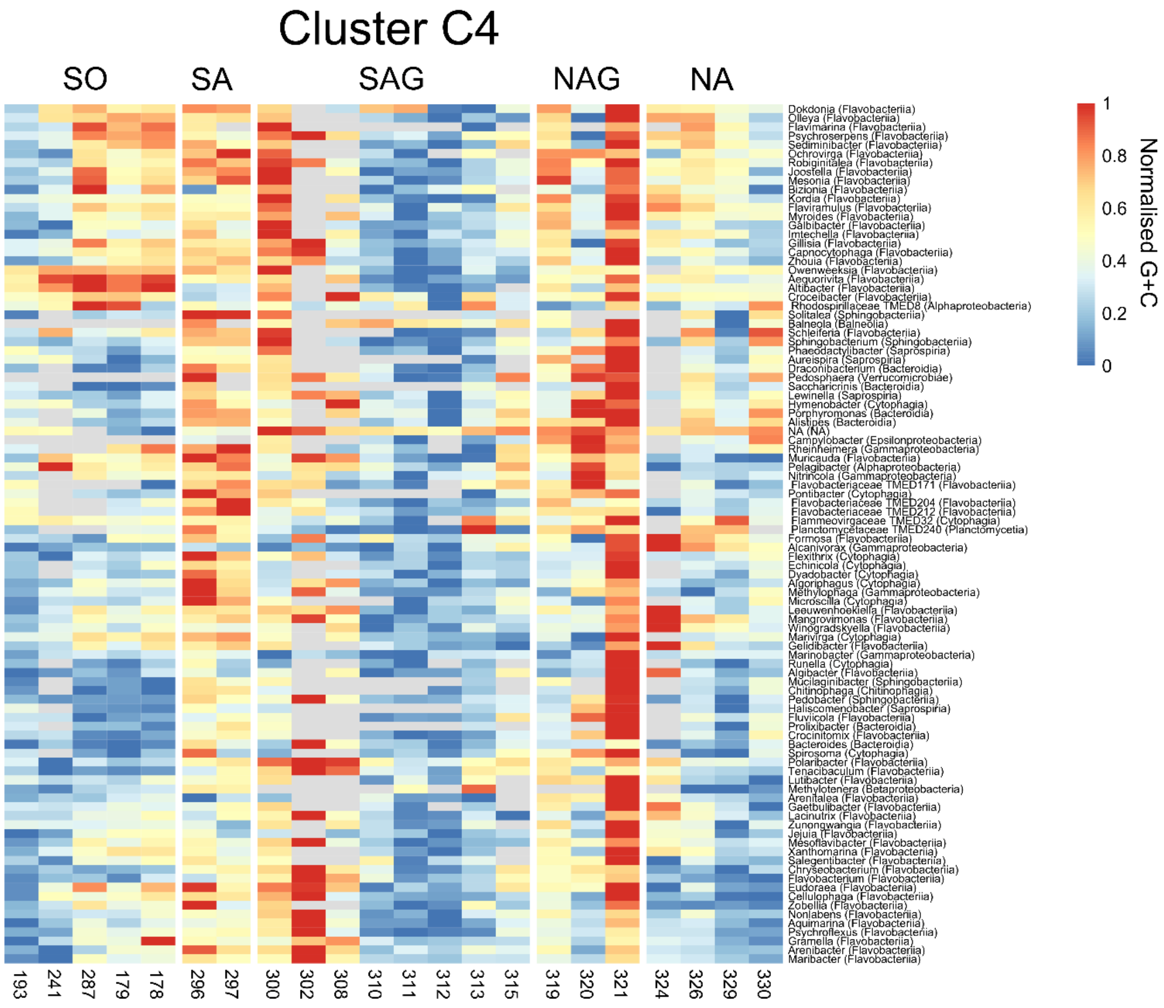

**Figure S5**

Normalized distribution of the G+C content of clusters C1 to C4 (Fig. S4) among prokaryotic and eukaryotic genera as well as viral families over the Atlantic Ocean transect between 62° and 47°N at stations 193 to 330. For station details see Table S1. Only taxa with ≥ 50 genes (~10kb) at ≥50% of stations were considered. The southernmost station 193, located in the Bransfield Strait with a limited dispersal rate[1], stood out and exhibited a lower AGL than the other stations in SO.

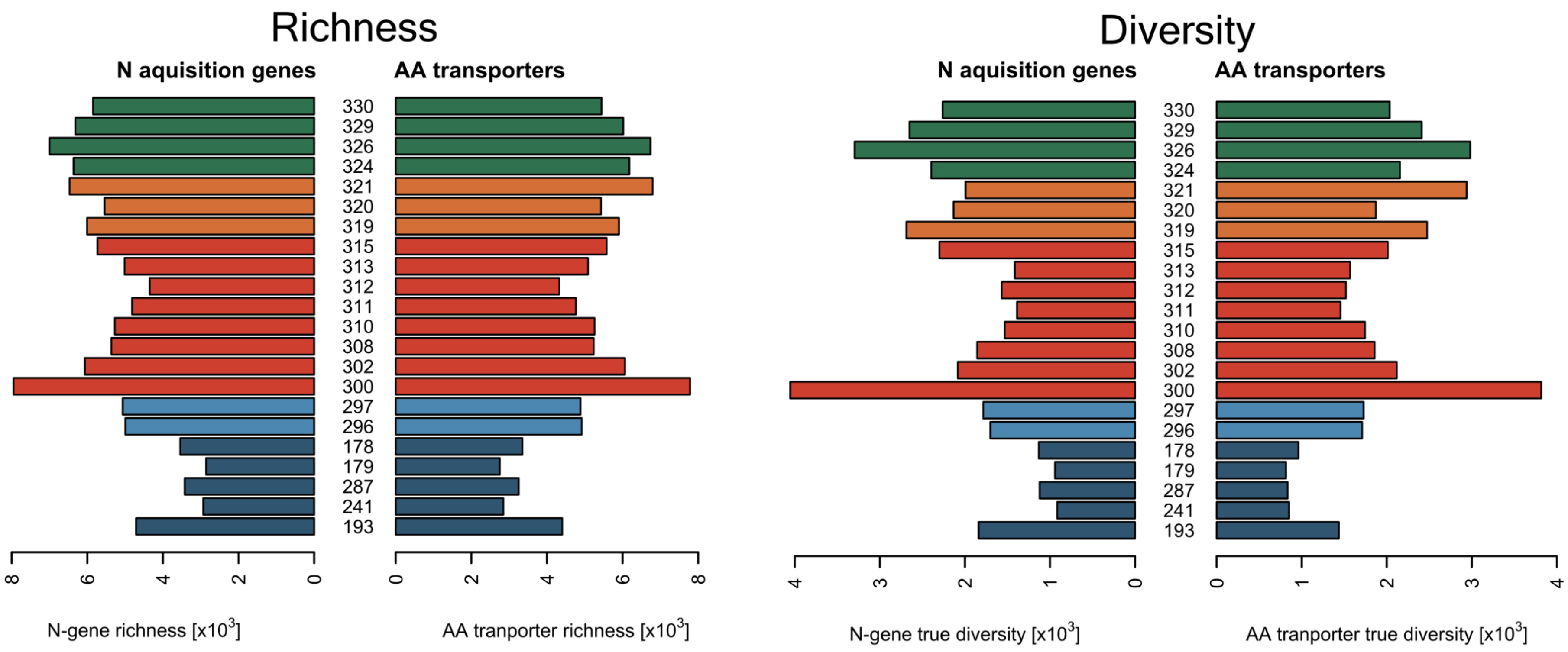

**Figure S6:**

Richness and Effective number of genes encoding N-acquisition and transport of amino acid (AA)-related N-compounds of the AOM at stations 193 to 330 along the Atlantic Ocean transect between 62° and 47°N.

.

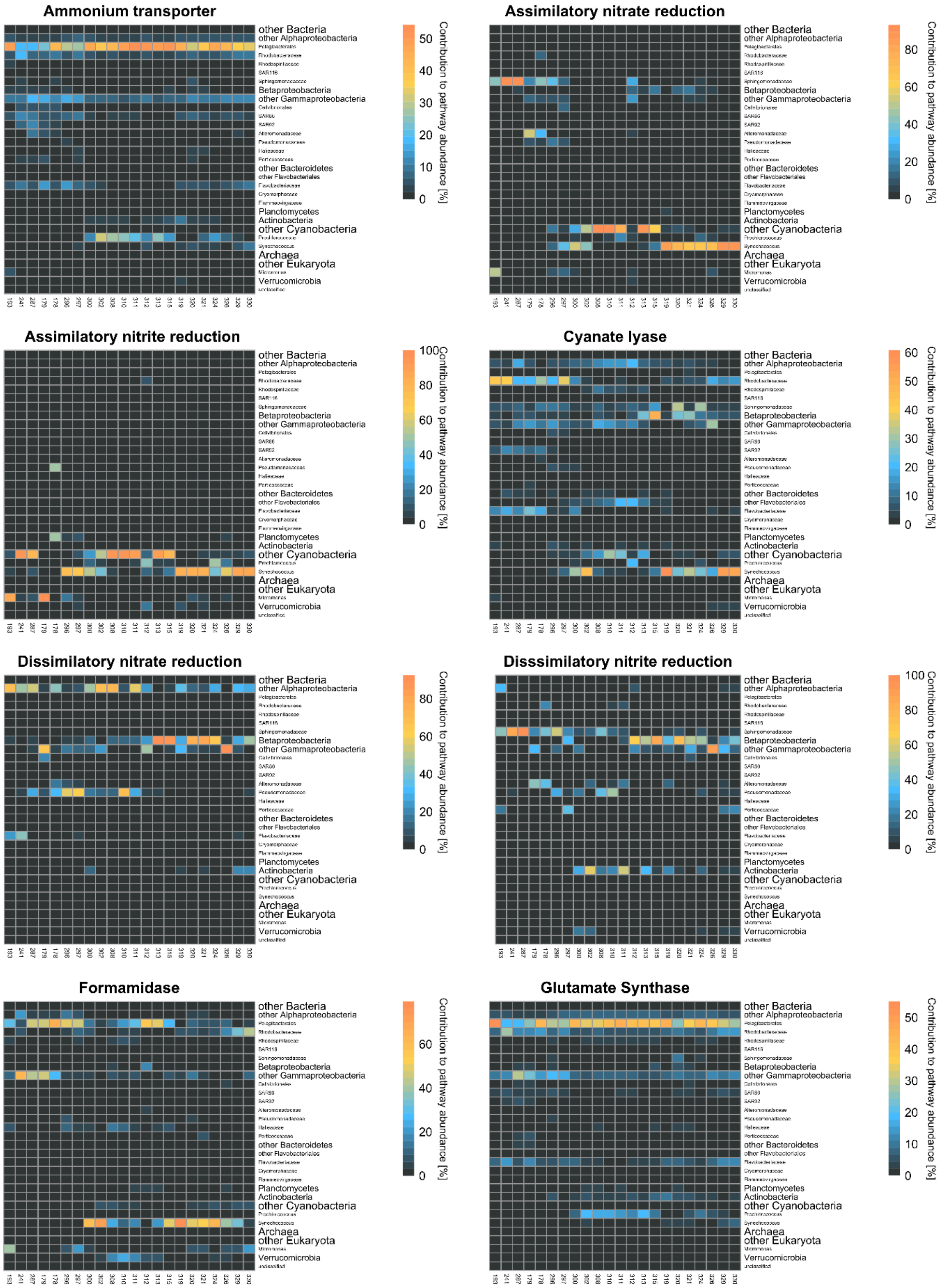
Ammonium transporter
Assimilatory nitrate reduction
Assimilatory nitrite reduction
Cyanate lyase
Dissimilatory nitrate reduction
Disssimilatory nitrite reduction
Formamidase
Glutamate Synthase
other Bacteria
other Alphaproteobacteria
Pelagibacterales
Rhodobacteraceae
Rhodospirillaceae
SAR116
Sphingomonadaceae
Betaproteobacteria
other Gammaproteobacteria
Cellvibrionales
SAR86
SAR92
Alteromonadaceae
Pseudomonadaceae
Halieaceae
Porticoccaceae
other Bacteroidetes
other Flavobacteriales
Flavobacteriaceae
Cryomorphaceae
Flammeovirgaceae
Planctomycetes
Actinobacteria
other Cyanobacteria
Prochlorococcus
Synechococcus
Archaea
other Eukaryota
Micromonas
Verrucomicrobia
unclassified
Contribution to pathway abundance [%]

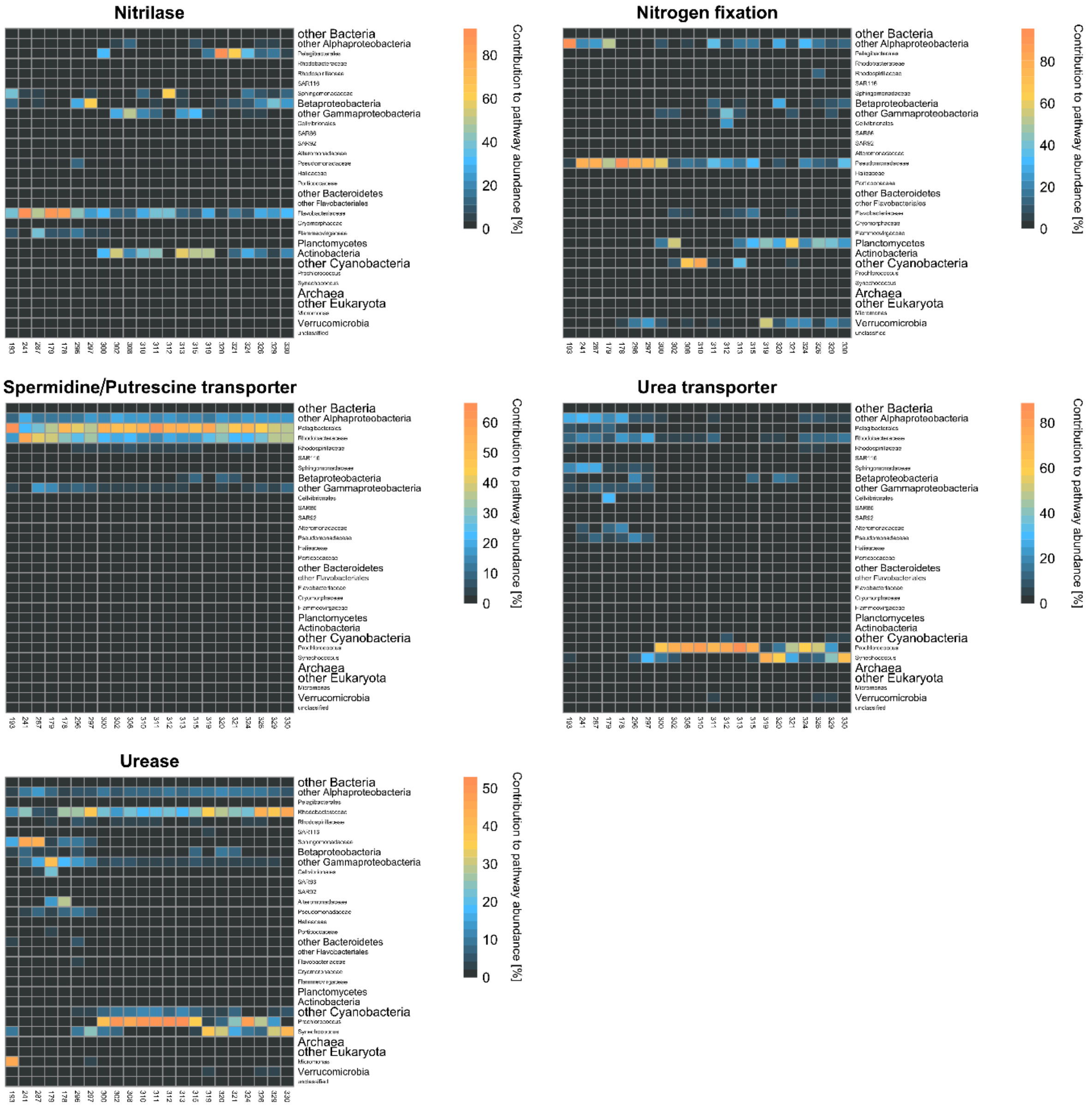

**Figure S7**

Distribution of various N-acquisition genes among prokaryotic families of the AOM at stations 193 and 330 along the Atlantic Ocean transect between 62°S and 47°N: assimilatory nitrate reduction, denitrification, cyanate lyase, ammonium transporter, urease, transporter of urea, nitrilase, oligo- and dipeptides, general and neutral amino acids, glycine betaine/proline and spermidine/putrescine.

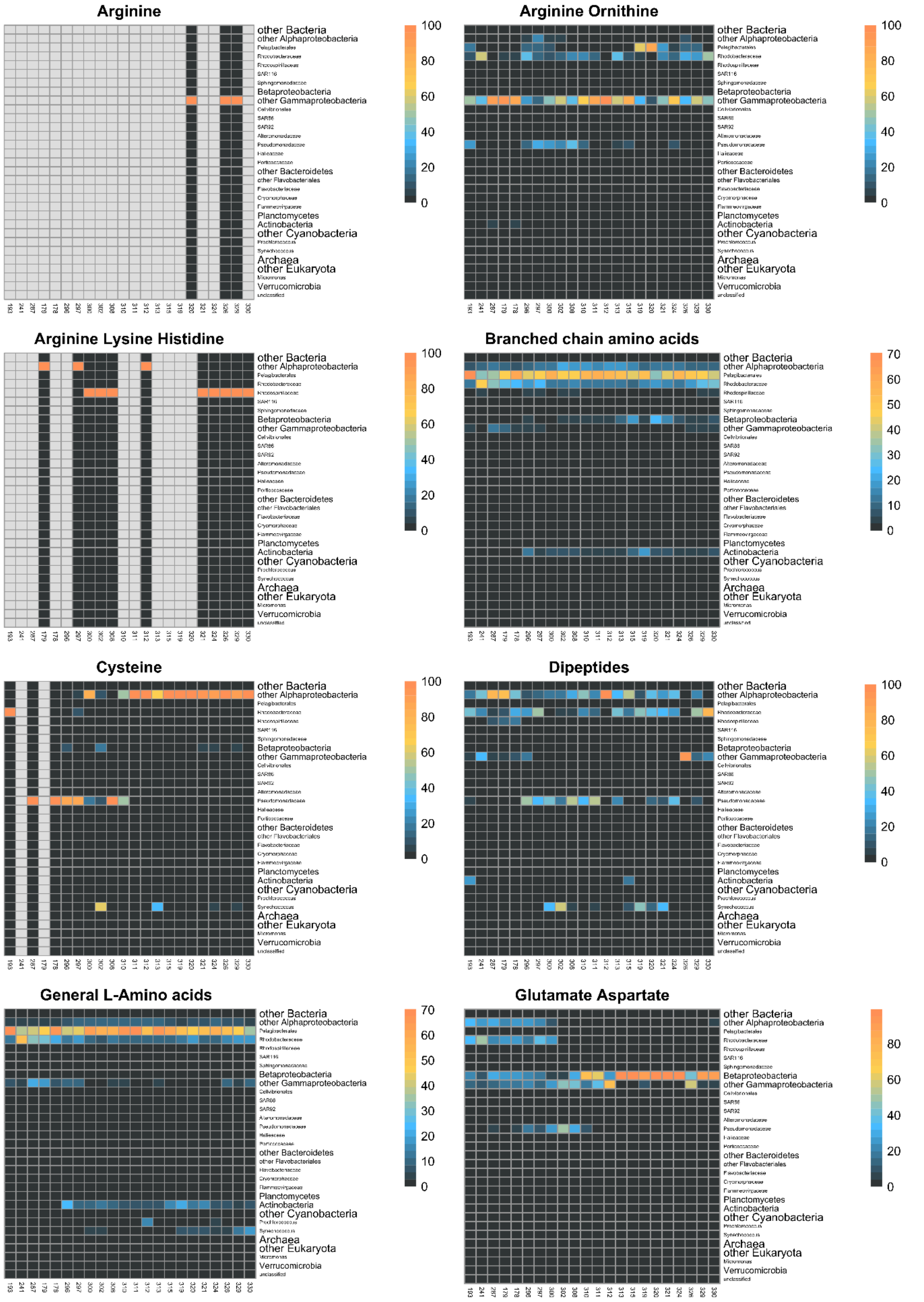
Arginine
Arginine Ornithine
Arginine Lysine Histidine
Branched chain amino acids
Cysteine
Dipeptides
General L-Amino acids
Glutamate Aspartate
other Bacteria
other Alphaproteobacteria
Betaproteobacteria
other Gammaproteobacteria
other Bacteroidetes
Planctomycetes
Actinobacteria
other Cyanobacteria
Archaea
other Eukaryota
Verrucomicrobia
100
80
60
40
20
0
70
50
30
10

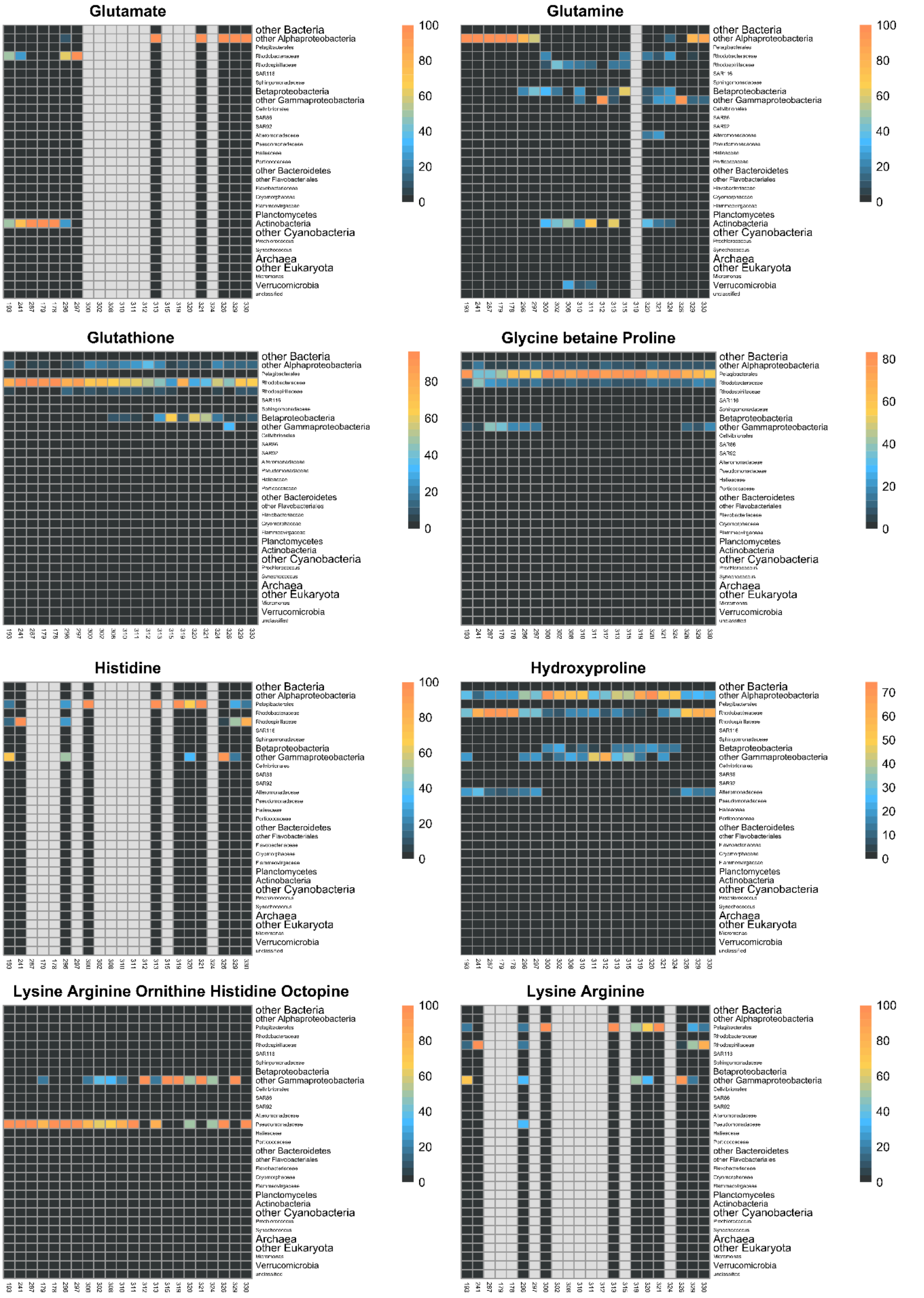
Glutamate
Glutamine
Glutathione
Glycine betaine Proline
Histidine
Hydroxyproline
Lysine Arginine Ornithine Histidine Octopine
Lysine Arginine
other Bacteria
other Alphaproteobacteria
Betaproteobacteria
other Gammaproteobacteria
other Bacteroidetes
Planctomycetes
Actinobacteria
other Cyanobacteria
Archaea
other Eukaryota
Verrucomicrobia
100
80
60
40
20
0
70
50
30
10

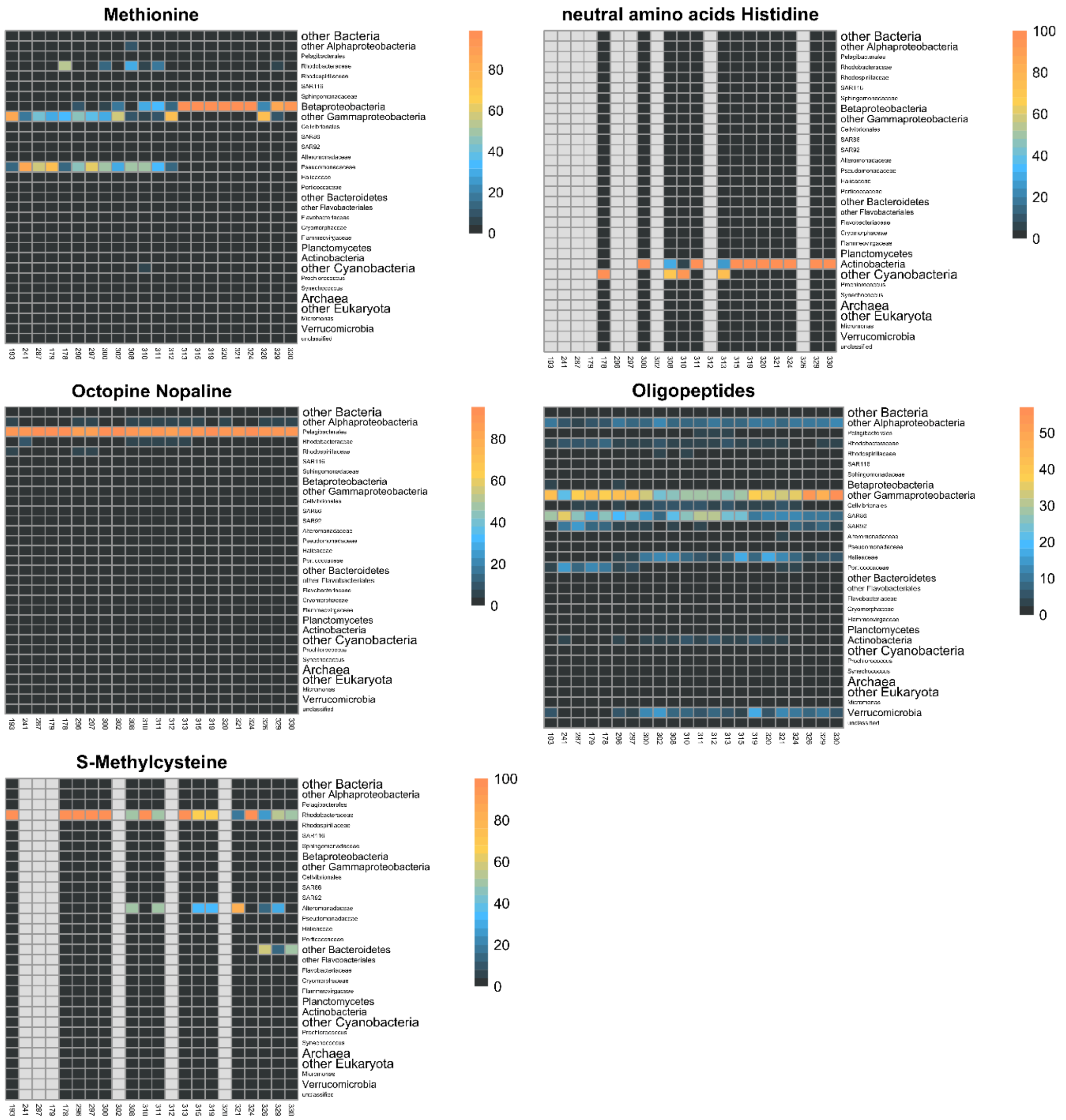

### Figure S8

Distribution of various amino acid transport systems among prokaryotic families of the AOM at stations 193 and 330 along the Atlantic Ocean transect between 62°S and 47°N.
